# Proteome-wide ubiquitinome profiling reveals substrate-specific dynamics within the USP7 network

**DOI:** 10.64898/2025.12.29.696849

**Authors:** Joyce Wolf van der Meer, Jan A. van der Knaap, Ayestha Sijm, Karel Bezstarosti, Dick H. W. Dekkers, Wouter A.S. Doff, Jeroen A. A. Demmers, C. Peter Verrijzer

## Abstract

USP7 is a pleiotropic deubiquitylating enzyme that is involved in tumor suppression, (neuro)development, chromatin regulation and the DNA damage response. How USP7 regulates these diverse pathways is still unclear. Here, we report data-independent acquisition and label free quantitation mass spectrometry (DIA-LFQ-MS) to profile the proteome-wide impact of USP7 on substrate de-ubiquitylation and overall protein abundance. First, we identified proteins associated with endogenous USP7 by immunopurification followed by DIA-LFQ-MS. Integration of our new results with earlier interactomes of epitope-tagged USP7 yielded a consensus set of high-confidence protein targets. Domain mapping analysis revealed that, in addition to the TRAF domain, the ubiquitin-like domains of USP7 play a key role in substrate selection. Using specific enrichment of tryptic K-ε-GG peptides, we mapped proteome-wide changes in ubiquitinome dynamics following inhibition of USP7. Combining unbiased proteome-wide and targeted quantitative mass spectrometry revealed that deubiquitylation by USP7 can have different effects on the stability of distinct substrates, and suggests that USP7’s activity profile is substrate-dependent rather than an intrinsic enzymatic property. Thus, in addition to providing a proteome-wide map of USP7 target sites, our multi-angle proteomics approach reveals that the effects of USP7-mediated deubiquitylation on its targets are remarkably variable and substrate-specific. Finally, based on these detailed molecular insights we show how USP7 connects various neurodevelopmental syndromes and tumor suppression pathways.

## INTRODUCTION

Ubiquitylation, the covalent attachment of the 76 amino acid (aa) ubiquitin (ub) polypeptide, regulates the stability and function of target proteins in eukaryotic cells (1–3). Ubiquitin signaling controls cellular processes ranging from proteostasis, cell division, protein localization, DNA repair, and gene expression. Consequently, dysregulation of cellular ubiquitin signaling plays major roles in a wide variety of human diseases (4). An E1-E2-E3 enzymatic cascade conjugates the carboxy-terminus of ub to (mostly) the ε-amino group of a lysine (K) residue in substrates (1–3). The formation of diverse forms of ub chains via ubiquitylation of one of its seven internal Ks, confers further functional information. E.g., K11 and K48-linked poly-ub chains mark proteins for proteasomal degradation, K63-linked chains assist the formation of large protein assemblies during DNA repair and mediate autophagic degradation, while mono-ubiquitylation of histone H2A on K119 (H2AK119ub1) marks Polycomb-repressed chromatin. Enzymatic cascades involving unique combinations of 2 E1, ∼40 E2 and ∼600 E3 ub ligases direct the ubiquitylation of thousands of cellular substrates (1–3, 5, 6). Ubiquitylation is reversed by selective deubiquitylation by ∼100 deubiquitylating enzymes (DUBs), providing an additional layer of regulation (7, 8). However, the molecular basis of substrate-selection and regulation by DUBs remains poorly understood.

Ubiquitin Specific Protease 7 (USP7) is an abundant nuclear DUB that has been implicated in tumor suppression, DNA repair, transcription and chromatin regulation (9–11). USP7 is essential for the development and viability of organisms ranging from *Drosophila* to mammals (12, 13). USP7 function is highly dosage-sensitive and haploinsufficiency of USP7 causes the neurodevelopmental disorder (NDD) Hao-Fountain Syndrome (OMIM 616863) (14–17). One of the most studied function of USP7 is its role in the MDM2-p53 tumor suppressor pathway (18, 19). Normally, USP7 functions as a negative regulator of p53 by deubiquitylating and stabilizing its E3 ubiquitin ligase MDM2. Under conditions of cellular stress, however, USP7 can directly deubiquitylate and stabilize p53 (19, 20). Co-deletion of *Tp53* gives only a partial rescue of the *Usp7*-null phenotype in mice (21), highlighting p53 independent functions of USP7. Indeed, USP7 is also part of the Polycomb gene repression system, and modulates the level of H2AK119ub1 through stabilization of the non-canonical Polycomb repressive complexes (ncPRC1s) ncPRC1.1 and ncPRC1.6 (12, 22–28). Recent research showed that epigenetic gene silencing by ncPRC1.1 is a major effector of USP7 function during neuronal differentiation (28). Additional USP7 targets include the epigenetic regulator PHIP/BRWD2 (25), the DNA repair factor ERCC6 (29, 30), the regulator of cytokinesis FBXO38 (31) and the RNA-helicase DHX40 (32). Notably, guanosine monophosphate synthase (GMPS) is a USP7-binding protein that regulates its enzymatic activity towards other substrates (12, 20, 33, 34). There has been a continuous accumulation of studies reporting novel USP7 substrates, albeit with varying degrees of confidence (35). Therefore, it is important to determine a core set of high-confidence substrates for this pleiotropic DUB.

The multidomain architecture of USP7 allows for intricate regulation and the ability to engage a plethora of substrates (9–11). The catalytic ub-hydrolase domain (CD) of USP7 is flanked by a N-terminal TRAF domain and five C-terminal ubiquitin-like domains (UBL1-5). The TRAF domain serves as a protein-protein interaction platform that mediates binding to multiple cellular interactors, including MDM2 and p53, but is also targeted by viral proteins such as EBNA1 and ICP0 (32, 36, 37). By itself, the CD has only very weak ub hydrolase activity. Full activity depends on activation by the unstructured 19 amino acids C-terminal peptide (CTP) of USP7 (38–41). UBL4-5 have been implicated in presenting the CTP at the catalytic site (9, 34). Additionally, the UBLs, in particular UBL1-2, can mediate binding to partner proteins (42).

Previous studies using unbiased MS methods to determine the USP7 interaction network mostly relied on over-expression of epitope-tagged USP7 (25, 26, 31, 32). We recently analyzed the interactome of endogenous USP7, immunopurified from SH-SY5Y neuroblastoma cells (28). Targeted quantitative MS was also used to accurately determine the effects of loss of USP7 or its inhibition on substrate abundance (25, 26, 28, 43). In a separate study, the impact of inhibition of USP7 on the ubiquitinome and proteome was determined using in-depth DIA-MS based ubiquitinomics, although these results were not integrated with USP7-binding data (44). Nevertheless, the identification of proteins displaying a rapid increase in ubiquitylation stoichiometry combined with reduction in overall protein abundance in this study, suggested that these are likely targets of USP7 (44). While, as expected, there is overlap between these independent studies, consensus high-confidence and integrated USP7 interactome and ubiquitinome profiles remains missing.

Here, we combined targeted proteomics and proteome-wide ubiquitinomics to determine the impact of USP7 on the ubiquitinome, the overall proteome and its direct substrates in a single study. For analysis of the dynamic ubiquitinome, we used the quantitative analysis pioneered by Steger and colleagues (44). Having established a high-confidence set of USP7 substrates, we provide a detailed proteome-wide map of USP7-regulated ubiquitylation sites. Pertinently, our results revealed unanticipated substrate-specific differences in deubiquitylation and protein stabilization dynamics of USP7. Finally, we discuss the implications of our findings for the role of USP7 in neurodevelopment and tumor suppression.

## EXPERIMENTAL PROCEDURES

### Experimental design and statistical rationale

The aim of this study was to comprehensively characterize 1) the USP7 interactome and the interactomes of a subset of binding partners of USP7 to define a detailed physical protein-protein interaction network, and 2) FT671-induced proteome and ubiquitinomics alterations and nominate candidate USP7 substrates by comparing control and FT671 treated cells using label-free quantitative DIA-MS. For all experiments in this manuscript three independent biological replicates were analyzed per condition. For each replicate, cells were independently cultured, harvested, lysed, digested and subsequently analyzed. Each LC–MS/MS run corresponds to a distinct biological sample. Samples were processed in parallel and randomized across the acquisition sequence with alternating control and IP or FT671-treated injections to minimize run-order and batch effects. A benchmark internal reference (HeLa digest) was injected at regular intervals to monitor instrument performance and signal stability over the course of the acquisition. DIA data were acquired using a variable window DIA method and analyzed with DIA-NN in library-based mode, using a project-specific spectral library or library-free (direct DIA) mode starting from a FASTA sequence database, in line with current guidelines for DIA-MS. Protein intensities from the two DIA-NN workflows were calculated using robust aggregation of tryptic peptide signals. Given the relatively small sample sizes in each experiment (3 *vs* 3), the study was designed as hypothesis-generating rather than fully powered for subtle quantitative differences. Benchmarking studies indicate that DIA-NN, in both library-free and library-based configurations, provides high quantitative precision and can detect large, consistent fold changes even at low *n*, although power to detect modest effects is limited. Differential protein or K-GG peptide abundance between control and IP, or control and FT671 treated samples was assessed using two-sided Welch’s t tests on LOG2-transformed protein intensities, which do not assume equal variances between groups. To account for multiple testing, resulting p values were adjusted using the Benjamini–Hochberg procedure, and both adjusted *p* values and effect sizes were used to prioritize candidates for follow-up validation. To focus on robust signals, proteins were required to be quantified in all replicates of at least one condition, supported by at least two proteotypic peptides in the DIA-NN reports, and to show a large effect size (absolute LOG2 fold change ≥ 1, i.e. ≥ 2-fold) together with *p* ≤ 0.05. For visualization (PCA, clustering, heatmaps), missing values were imputed using a left-censored approach suitable for DIA LFQ data, whereas all hypothesis testing was performed on non-imputed intensities after filtering to avoid artefactual significance due to imputation.

### Materials

For materials see Supplemental Table S1.

### Cloning procedures

All PCR reactions for the production of constructs were performed using KOD-hotstart polymerase. RNA was isolated from HEK293T or U2OS cells using TRI reagent according to the manufacturer’s instructions, followed by cDNA synthesis using SuperScript II Reverse Transcriptase. RADX, TMPO, PJA1, and RNF220 were amplified from HEK293T cDNA, while MAGED4 and PJA2 were produced from U2OS cDNA. pENTR11-3xFlag-USP7 (Sijm et al., 2022) was used as PCR template for the generation of USP7 deletion constructs. Amplified PCR products were cloned into pENTR11 (Invitrogen) or pGEX2TK for expression in *E.coli*. The cDNA’s of pENTR11 constructs were recombined into pDEST-3xFLAG or a gateway compatible version of pQCXIP (pQCXIP-DEST) using LR Clonase II according to the manufacturer’s instructions. All plasmids were produced in *E.coli* TOP10. The integrity of all constructs was verified by Sanger sequencing of the coding sequences and flanking regions. See Supplemental Table S1 for an overview of all oligonucleotides used for cloning.

### Tissue culture

All cells were maintained in a humidified incubator with 5% CO_2_ at 37°C. U2OS, HEK293T cells and HEK293T USP7-KO cells were cultured in Dulbecco’s modified Eagle’s medium (DMEM; Gibco) supplemented with 10% fetal bovine serum (FBS) and 1x Pen/Strep (100 units/ml Penicillin and 100 µg/ml Streptomycin). Cell cultures were split at 80-90% confluency by aspiration of the media, one wash with PBS (137 mM NaCl, 2.7 mM KCl, 10 mM Na_2_HPO_4_ and 1.8 mM KH_2_PO_4_, pH 7.4) and subsequent trypsinization by incubation with PBS containing 0.05% Trypsin and 0.53 mM EDTA. After detachment, cells were resuspended in medium containing 10% FBS and 1x Pen/Strep, and split. All cultures were routinely checked for mycoplasma using the MycoAlert kit.

### GST-pull-down assays

GST-fusion proteins were expressed and affinity purified(AP) essentially as described in (45). In short, *E.coli* BL21-CodonPlus(DE3)RIL strains containing the appropriate expression plasmid were grown at 37°C in LB medium containing 100 µg/ml ampicillin under constant shaking until OD_600_ 0.4-0.8 was reached. Expression of the GST-tagged proteins was induced by addition of 0.1 mM Isopropyl β-D-1-thiogalactopyranoside (IPTG, Sigma Aldrich 367-93-1), followed by an additional incubation at 25°C for 2 hours (h). Upon harvesting and lysis, GST-fusion proteins were purified from the soluble fraction using Glutathione Sepharose® 4 Fast Flow resin, using standard procedures (45). Next, 500 mg GST-tagged protein was cross-linked to the Glutathione Sepharose Fast Flow resin using dimethylpimelimidate dihydrochloride. Whole cell extracts (WCE) were prepared from two 15 cm dishes of HEK293T, which were harvested by scraping, washed with PBS and collected by centrifugation in a precooled centrifuge at 4°C and 1000 g. All following steps were carried out on ice or at 4°C. Cell pellets were resuspended in 1ml NEH buffer (50mM HEPES-KOH pH 7.6, 150 mM NaCl, 5 mM EDTA, 1.5 mM MgCl_2_, 0.1% Nonidet P40, 10% glycerol, 1mM DTT and protease inhibitors: 1µg/ml pepstatin, 1µg/ml aprotinin, 1µg/ml leupeptin, 0.2 mM AEBSF) and sonicated for 5 minutes (min) in a Bioruptor UCD-200 sonicator using a 30 sec on/off cycle. Protein concentration of the WCE was determined by Bradford assay. WCE was stored by snap freezing in liquid nitrogen and stored at −80°C. After thawing the WCE was cleared by centrifugation for 15 min at 16000 g. To determine binding to the recombinant USP7 domains, 20 μl of GST-USP7 fusion domain crosslinked to Glutathione Sepharose was washed three times with HEMG/150/0.1% Nonidet P-40 (25 mM HEPES KOH pH7.6, 0.1 mM EDTA, 12.5 mM MgCl_2_, 10% glycerol, containing 150 mM KCl, 0.1% Nonidet P40, and protease inhibitors: 1µg/ml pepstatin, 1µg/ml aprotinin, 1µg/ml leupeptin, 0.2 mM AEBSF) and incubated with 1 mg WCE for 2 h on a rotating wheel. Next, the resin was washed five times using HEMG/150/0.1% Nonidet P-40, followed by three washes with HEMG/150/0.01% Nonidet P-40 and one quick wash with HEMG/100 w/o AEBSF. All washes were performed by addition of 1ml of buffer, incubation for 10 min on a rotating wheel and centrifugation at 800 g for 3 min, followed by removal of the supernatant. The resin was stored in HEMG/100 w/o AEBSF and bound proteins were analyzed by mass spectrometry.

### Immunopurification of Flag-tagged proteins

Immunopurifications of Flag-tagged protein were performed in three biological replicates and essentially performed as described in (26). Briefly, per replicate two 15 cm dishes of HEK293T cells were transfected with the appropriate expression vectors (20 µg plasmid DNA per plate) using polyethylenimine (PEI). After 48 h cells were scraped in PBS, and collected by centrifugation in a precooled centrifuge at 4°C, 1000 g for 5 min. Expression of the Flag-tagged proteins was verified by western blotting using anti-Flag antibodies. All following steps were on ice or at 4°C. To purify Flag-tagged proteins, cell pellets were resuspended in 1 ml NEH buffer w/o DTT and sonicated in a Bioruptor UCD-200 sonicator for 5 min using a 30 sec on/off cycle. Cell lysates were incubated with 100 U benzonase (Novagen) for 30 min on the rotating wheel. Ethidium bromide was added to a final concentration of 50 µg/ml, lysates were cleared by centrifugation at 20,000 g for 20 min and the supernatants were transferred to new 1.5 ml Eppendorf tubes. Anti-Flag M2 affinity gel was prepared by washing three times with PBS, once with 0.1 M glycine pH 3.5 and 150mM NaCl and again three times with PBS. The affinity gel was left in PBS over night at 4° C to renature. Next day, the anti-Flag M2 affinity gel was equilibrated with NEH buffer w/o DTT and 20 µl of gel was added to each of the prepared cell lysates and incubated for 2.5 h on a rotating wheel. Next, the anti-Flag affinity gel was washed 7 times with HEG/150/0.1% Nonidet P-40, (50 mM HEPES KOH pH7.6, 0.1 mM EDTA, 10% glycerol, containing 150 mM NaCl, 0.1 % Nonidet P40 and a cocktail of protease inhibitors), two times with HEG/150/0.01% Nonidet P-40 and finally 2 times with HEG/100 w/o AEBSF. The resulting immunopurified proteins bound to the gel were directly processed for mass spectrometry.

### Immunopurification of endogenous USP7

Each purification was performed using biological triplicates. HEK293T or HEK293T USP7-KO cells were harvested from two 90% confluent 15 cm plates per triplicate by scraping in PBS. Following centrifugation at 1000g for 5 min, cell pellets were lysed in 1 ml ice cold HMG/150 (25 mM HEPES pH 7.6, 2.5 mM MgCl_2_, 10% glycerol, 0.5% Nonidet P-40, containing 150 mM KCl and protease inhibitors: 1µg/ml pepstatin, 1µg/ml aprotinin, 1µg/ml leupeptin, 0.2 mM AEBSF). The lysates were sonicated for 5 min in a Bioruptor UCD-200 sonicator using 30 sec ON/OFF cycles and subsequently incubated with 100 U benzonase for 30 min at 4°C. Digestion was stopped by the addition of 5 µl 0.5 M EDTA pH 8.0. Next, lysates were transferred to 1.5 ml eppendorf tubes, snap frozen in liquid N_2_ and stored at −80°C. All subsequent steps were performed at 4°C. Extracts were thawed on ice, centrifuged at 16000 g for 25 min and supernatants were transferred into low protein binding Eppendorf tubes. Each cleared lysate was incubated with 22.5 µl anti-USP7 antibody (Bethyl Laboratories, # A300-033A) on a rotating wheel for 2 h. Next, 15 µl of equilibrated protein A Sepharose (bead volume) was added to each tube and extracts were incubated on a rotating wheel for an additional 2 h. Following centrifugation at 800 g for 3 min, the resin was washed three times with 1 ml HEMG/150/0.1% Nonidet P40, three times with HEMG/150/0.01% Nonidet P40, once with HEMG/100/0.01% Nonidet P40, and finally twice with HEMG/100, w/o AEBSF). The resulting immunopurified proteins bound to the resin were analyzed by mass spectrometry.

### Sample preparation for mass spectrometry

For the analysis of immunopurified samples, proteins were digested on-bead with sequencing grade trypsin (1:100 (w:w), Roche) overnight at room temperature. Protein digests were desalted using a C18 Stagetip (2 plugs of 3M Empore C18) and eluted with 80 % acetonitrile and dried in a Speedvac centrifuge. Peptides were then analyzed by nanoflow LC-MS/MS as described (28). For **global proteome analysis** of whole cell extracts, cells were lysed in 100 mM Tris/HCl, pH 8.2, containing 1 % sodium deoxycholate (SDC) using sonication in a Bioruptor Pico (Diagenode). Protein concentrations were measured using the BCA assay (ThermoFisher Scientific). 100 μg protein was reduced in lysis buffer with 5 mM dithiothreitol and alkylated with 10 mM iodoacetamide. Next, proteins were digested with 2.5 μg trypsin (1:40 enzyme:substrate ratio) overnight at 37 °C. After digestion, peptides were acidified with trifluoroacetic acid (TFA) to a final concentration of 0.5 % and centrifuged at 10,000 g for 10 min to spin down the precipitated SDC. Peptides in the supernatant were desalted on a 50 mg C18 Sep-Pak Vac cartridge (Waters). After washing the cartridge with 0.1 % TFA, peptides were eluted with 50 % acetonitrile and dried in a Speedvac centrifuge. Peptides were then analyzed by nanoflow LC-MS/MS as described below. For **ubiquitinome analysis**, biological triplicates of HEK293T cells were grown in 15 cm plates till 70% confluency and treated with 10 µM FT671 (1 and 6 h) or its solvent, 7 mM DMSO (6 h). Cells were washed twice with PBS and subsequently processed for mass spectrometry. Ubiquitin remnant motif (K-ε-GG) antibodies coupled to beads (PTMscan, Cell Signaling Technologies) were used for selective enrichment of tryptic K-GG peptides according to the manufacturer’s protocol. A batch of beads was washed twice with PBS and peptide fractions were dissolved in 1.4 ml IAP buffer (50 mM MOPS, 10 mM sodium phosphate and 50 mM NaCl, pH 7.2) and debris was spun down. The supernatant was incubated with beads for 2 h at 4 °C on a rotator unit. Beads were transferred into a P200 pipette tip equipped with a GF/F filter plug (Whatman part #1825-021) to retain the beads. The beads were then washed 3 times with 200 μl of ice-cold IAP buffer and subsequently 5 times with 200 μl of ice-cold milliQ H_2_O and peptides were eluted using 2 cycles of 50 μl 0.15% TFA. Finally, peptides were desalted using a C18 stage tip and dried to completeness using vacuum centrifugation. K-GG peptides were then analyzed by nanoflow LC-MS/MS as described below.

### Mass spectrometry

**Nanoflow LC-MS/MS** was performed on an EASY-nLC system (Thermo) coupled to an Orbitrap Eclipse Tribrid mass spectrometer (both ThermoFisher Scientific) or on a Vanquish Neo LC system (Thermo) coupled to an Orbitrap Exploris 480 (Thermo), all operating in positive mode and equipped with a nanospray source. Peptide mixtures were trapped on a PepMap trapping column (2 cm × 100 µm, Thermo, 164750) at a flow rate of 1 µl/min. Peptide separation was performed on ReproSil C18 reversed phase column (Dr Maisch GmbH; column dimensions 25 cm × 75 µm, packed in-house) using a linear gradient from 0 to 80% B (A = 0.1% FA; B = 80% (v/v) AcN, 0.1 % FA) in 120 min and at a constant flow rate of 250 nl/min. The column eluent was directly sprayed into the ESI source of the mass spectrometer. For **data dependent acquisition** (DDA): All mass spectra were acquired in profile mode and the resolution in MS1 mode was set to 120,000 (automatic gain control (AGC) target: 4E5) and the m/z range to 350-1400. Fragmentation of precursors was performed in 2 s cycle time data-dependent mode by higher-energy collisional dissociation (HCD, or beam-type collision induced dissociation (CID)) with a precursor window of 1.6 m/z and a normalized collision energy of 30.0; MS2 spectra were recorded in the orbitrap at 30,000 resolution. Singly charged precursors were excluded from fragmentation and the dynamic exclusion was set to 60 seconds. For **data independent acquisition** (DIA): All spectra were recorded at a resolution of 120,000 for full scans in the scan range from 350–1100 m/z. The maximum injection time was set to 50 ms (AGC target: 4E5). For MS2 acquisition, the mass range was set to 336–1391 m/z with variable isolation windows ranging from 7–82 m/z with a window overlap of 1 m/z. The orbitrap resolution for MS2 scans was set to 30,000. The maximum injection time was at 54 ms (AGC target: 5E4; normalized AGC target: 100 %). For **targeted proteomics**, a PRM regime was used to select for a set of previously selected peptides on an Orbitrap Eclipse Tribrid or an Orbitrap Exploris 480 mass spectrometer operating in positive mode. Precursors were selected in the quadrupole with an isolation width of 0.7 m/z and fragmented with HCD using 30 % collision energy (CE). MS1 and MS2 spectra were recorded in the orbitrap at 30,000 resolution in profile mode and with standard AGC target settings. The injection time mode was set to dynamic with a minimum of 9 points across the peak. The sequence of sampling was blanks first and then in order of increasing peptide input amounts to avoid any contamination of previous samples.

### Data analysis

**DDA raw data** files were analyzed using the MaxQuant software suite (version 2.2.0.0, www.maxquant.org, (46) for the identification and relative quantification of proteins. ‘Match between runs’ was disabled and a false discovery rate (FDR) of 0.01 for peptides and proteins and a minimum peptide length of 6 amino acids were required. The Andromeda search engine was used to search the MS/MS spectra against the *Homo sapiens* Uniprot database (version May 2022) concatenated with the reversed versions of all sequences and a contaminant database listing typical background proteins. A maximum of two missed cleavages were allowed. MS/MS spectra were analyzed using MaxQuant’s default settings for Orbitrap and ion trap spectra. The maximum precursor ion charge state used for searching was 7 and the enzyme specificity was set to trypsin. Further modifications were cysteine carbamidomethylation (fixed) as well as methionine oxidation (variable). The minimum number of peptides for positive protein identification was set to 2. The minimum number of razor and unique peptides set to 1. Only unique and razor non-modified, methionine oxidized and protein N-terminal acetylated peptides were used for protein quantitation. The minimal score for modified peptides was set to 40 (default value). **DIA raw data files** were analyzed with DIANN (versions 1.9.1 through 2.2.0 were used http://www.github.com/vdemichev/DiaNN), with standard settings, including N-terminal Met excision and Cys carbamidomethylation. In addition, the K-GG and Ox(M) options was checked and the number of missed cleavages was raised to 2. The same fasta reference database was used as for the DDA MaxQuant searches. **DIANN output files** were loaded into Perseus. Only identified proteins with at least three valid values in at least one group were included. After grouping replicates a two-sample t test was performed. For truncation, Benjamini-Hochberg FDR was applied. For profile plots, Z scoring was applied. **PRM data** were analyzed with Skyline (version 23.0.1.268, http://skyline.ms). Skyline reports containing all essential data such as area per fragment for each target were loaded into R and manually curated. If the expected peptide fragment ion peaks could not be correctly assigned, fragments were excluded and their associated areas were set to 0. Summed areas were calculated for all fragment ion peak chromatograms for each target peptide and then summed over all targeted peptides per protein. Standard deviations of at least three biological replicates were calculated for every target peptide in each condition and indicated in the bar plots. All summed areas were normalized to the WT condition. Peptide bar graphs were constructed by dividing the bar value for each target peptide into the respective areas for every fragment ion peak. For the protein abundance comparisons, the normalized areas for all detected fragment ion chromatograms were summed over all selected target peptides per protein and visualized in box plots. The median for the WT condition was set to 1. All protein abundances are based on ≥ 2 target tryptic peptides unless indicated otherwise. **Downstream data analysis** was performed using Perseus (www.maxquant.org/perseus/), Prism (v 10, www.graphpad.com) or in-house developed software tools. In Perseus, data from MaxQuant or DIANN searches were imported, and all peptide and protein intensity data were first LOG2 transformed. For the analysis of putative interactors in IP analyses, and the identification of up- or downregulated target proteins and their ubiquitination sites upon USP7i treatment, Student’s double-sided **t testing** was performed on LOG2 DIA based LFQ intensity data of triplicate experiments, with permutation-based or Benjamini-Hochberg FDR (0.05) truncation (250 randomizations). MS raw data and data for protein identification and quantification were submitted as Supplemental tables to the **ProteomeXchange Consortium** via the PRIDE partner repository with the data identifier PXD071773.

### Analysis of independent MS results

To derive a consensus set of USP7-binding proteins (Fig. 1*B*), we compared results for endogenous USP7 from the present study (Fig. 1*A*) with four previously reported IP-MS datasets from our own and other labs (25, 26, 28, 31). For (31), interactors were included if the peptides number in the Flag-USP7 IP from AGS cells was at least 10-fold higher than that in the bGAL control IP. For (25), the SFB-USP7 binders in HEK293A from supplemental data S5 were used. For the endogenous USP7-IP in HEK293T cells (this study) and SH-SY5Y (28), interactors of endogenous USP7 in SHSY5Y cells were determined by taking the average pg-Quantity values and selecting hits that have a >10-fold USP7-IPs/control average pg-quantity and are present in all three independent IPs. Interactors of Flag-USP7 in HEK293T cells (26) were selected by taking the average iBAQ values and selecting hits that have a >10-fold USP7-IPs/control average iBAQ value, and are present in all three IPs.

**Fig. 1.**
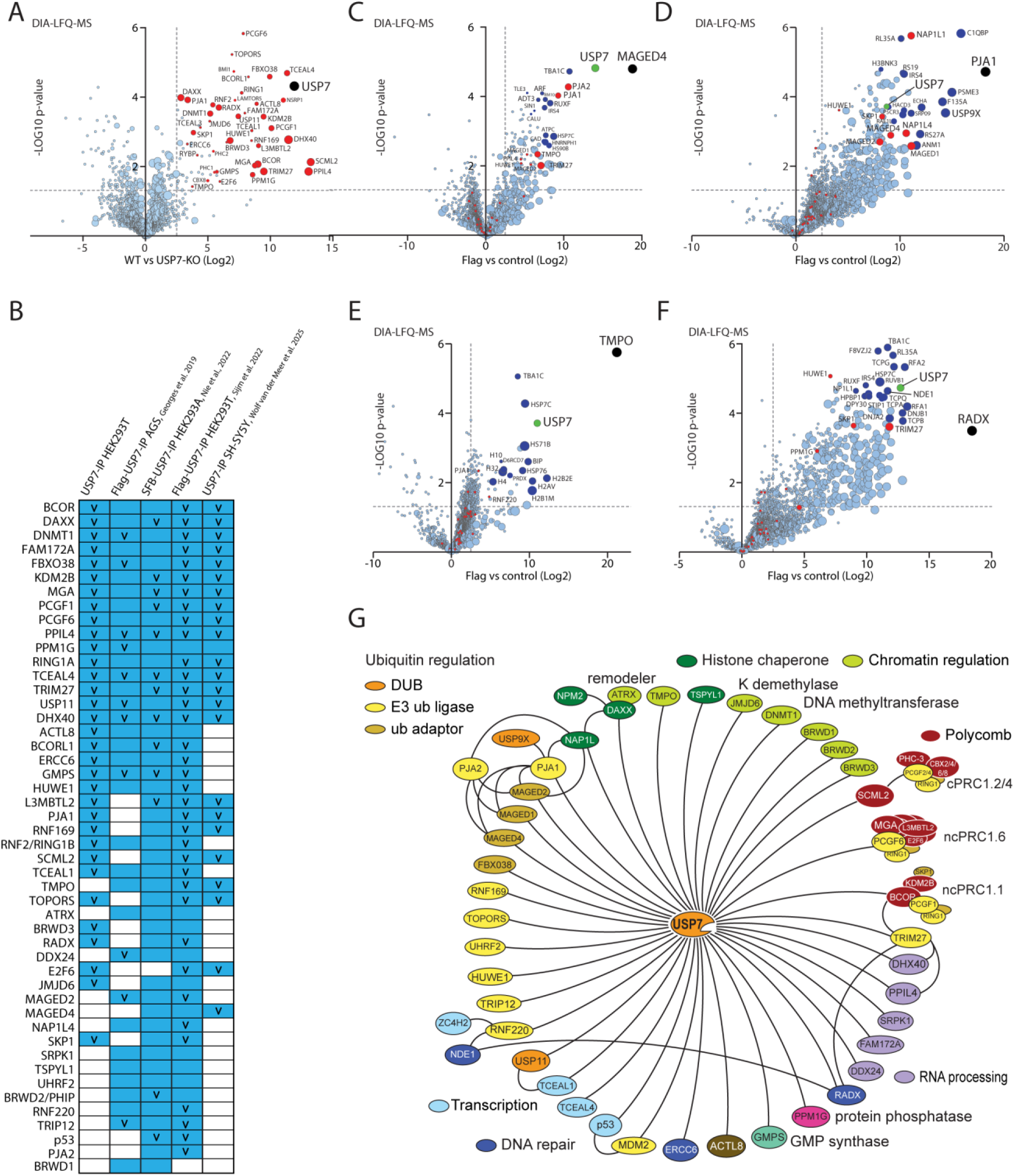
Establishment of a high confidence USP7 interactome. ***A***, proteins associated with endogenous USP7 purified from HEK293T cells. Volcano plot representing a statistical double-sided t-test based on LFQ quantitative MS data of protein enrichment in IPs from WT versus USP7-KO cells. The size of the data spheres reflects the relative abundance of the protein in the IP analysis, based on its mass spectrometry LFQ intensity. The dashed grey lines indicate arbitrary thresholds (i.e., p value = 0.05, LOG2 t test difference = 2 or enrichment over control >4-fold). The analysis is based on three independent biological replicate experiments. High-confidence USP7-interacting proteins are shown in red. The bait (USP7) is indicated in black. For the complete data set see Supplemental Table S2. ***B***, Consensus set of high confidence USP7-associated proteins. Significant USP7-associated proteins (blue) identified by IP-MS analysis of endogenous USP7 from HEK293T cells (present study), Flag-USP7 from AGS gastric cancer cells (31), SFB-USP7 from HEK293A cells (25), Flag-USP7 from HEK293T cells (26), and endogenous USP7 from SH-SY5Y cells (28). “v” denotes proteins that were marked as USP7-interacting protein in one of the previous studies. ***C*-*F***, IP-MS analysis of Flag-MAGED4, Flag-PJA1, Flag-TMPO and Flag-RADX expressed in HEK293T cells. Analysis as described in (A), LFQ quantitative MS data of protein enrichment in IPs from cells expressing a Flag-tagged bait is plotted versus IPs from cells that do not. Baits are indicated in black, USP7 in green, USP7-interacting proteins are shown in red and other interacting factors in blue. For the complete data set see Supplemental Table S2. *G*, A schematic summary of USP7-centered interaction network. PRC1 subunits are depicted as part of established complexes. Main molecular functions are indicated.

## RESULTS

### Determination of the high-confidence USP7 interactome

We extended previous USP7 interaction studies (25, 26, 28, 31, 32) by immunopurification (IP) of endogenous USP7 from whole cell extracts (WCEs) of HEK293T human embryonic kidney cells, followed by protein identification by data-independent acquisition label-free quantification mass spectrometry (DIA-LFQ-MS). As a negative control, we used anti-USP7 IPs from WCEs of cells in which we had deleted the *USP7* gene by CRISPR-Cas9 genome editing (HEK293T USP7-KO). All IPs were performed in three biological replicate experiments, and followed by on-bead digestion. Tryptic peptides were identified by nanoflow liquid chromatography (nLC) followed by DIA-LFQ-MS. To identify significant interacting proteins, we performed a statistical double-sided t-test based on LFQ quantitative MS data of protein enrichment in IPs from WT versus USP7-KO cells (Fig. 1*A* and Supplemental Table S2). Additionally, the protein abundance in the IPs based on the summed mass spectral peptide intensities for each protein hit (arbitrary units) is reflected by the size of the data spheres in the plots. This analysis confirmed the presence of generally accepted USP7 partners, including DHX40, DNMT1, FBXO38, ncPRC1.1, ncPRC1.6, PHIP, TCEAL4 and TRIM27, but also identified novel or less-established associated proteins, such as ACTL8, BRWD3, FAM172A, JMJD6, PJA1, RADX, RNF169 and TOPORS.

To derive a consensus set of USP7-binding proteins, we compared our results with four previously reported IP-MS datasets from our own and other laboratories. We juxtaposed the IP-MS results from endogenous USP7 isolated from HEK293T cells (this study), Flag epitope-tagged USP7 (Flag-USP7) from AGS gastric cancer cells (31), S-protein-FLAG-Streptavidin binding tagged USP7 (SFB-USP7) from HEK293A cells (25), Flag-USP7 from HEK293T cells (26) and endogenous USP7 IPed from SH-SY5Y neuroblastoma cells (28). We note that, due to the expression of viral oncoproteins, the p53-MDM2 pathway is disrupted in HEK293T and HEK293A cells. Next, we compared the MS datasets from these publications to evaluate the presence and enrichment of proteins that may not have been named in the associated publication (see methods and legends for detailed selection criteria). This comparison yielded a set of high-confidence USP7-associated proteins of which 16 were enriched in all five independent experiments, 13 in four and 16 in three out of five independent IP-MS studies (Fig. 1*B*). Proteins that were specifically mentioned in one of these previous studies are indicated by a “v”. Earlier MS studies revealed that USP7-binding proteins MGA, PCGF6, L3MBTL2, BCOR, PCGF1, KDM2B, USP11, RNF220, BRWD2/PHIP, and TRIM27 showed reduced levels in USP7-KO cells or following long-term inhibition of USP7 (25, 26, 28).

To explore the USP7 network further, we previously used IP-MS of Flag-tagged BCOR, DAXX, GMPS, MAGED2, MGA, PCGF1, PCGF6, SCML2, TRIM27 and USP11 to confirm their association with USP7 (26). We now extended our analysis by IP-MS of Flag-tagged MAGED4, PJA1, TMPO and RADX (Fig. 1*C*-*F* and Supplemental Table S2). These reciprocal IP-MS experiments confirmed the association of all four proteins with USP7. Notably, only few of the other USP7-interacting factors (indicated in red) showed significant enrichment in the reciprocal IP-MS plots, emphasizing that USP7 is at the center of a multi-nodal network of distinct hubs. The IP-MS analysis of RADX revealed binding to TRIM27, RPA and NDE1. Conversely, RADX was also identified in IP-MS analysis of TRIM27 (26). IP-MS analysis of PJA1, PJA2, MAGED2 (26) and MAGED4 revealed the association of PJA1 with MAGED1, MAGED2, MAGED4, USP9x, NAP1L1/4 and USP7. We also performed an IP-MS analysis of Flag-tagged PJA2 but observed no significant binding to USP7 (Supplemental Fig. S1*A* and Supplemental Table S2). Instead, PJA2 was found associated with MAGED1, MAGED2, MAGED4, USP9x and NAP1L1/4. Thus, PJA1 and PJA2 are part of partially overlapping, yet distinct protein networks. We suspected that RNF220 might be a USP7 substrate because we detected it previously in USP7 IPs (26), and its abundance is highly sensitive to USP7 inhibition (26, 43, 44). However, although IP-MS analysis of Flag-tagged RNF220 identified a stable association with ZC4H2 and NDE1, it did not reveal significant binding to USP7 (Supplemental Fig. S1*B* and Supplemental Table S2). Thus, a potential association between RNF220 and USP7 is likely to be transient. Our current view of the USP7 interaction network is summarized in Fig. 1*G*.

In conclusion, our IP-MS analysis of endogenous USP7, combined with additional and reciprocal IP-MS studies from our own and other labs, yielded a high-confidence USP7 network. Further analysis revealed that this extensive network comprises a variety of USP7-hubs, that are largely isolated from each other and dedicated to distinct cellular functions. Although transient interactions or cell type specific associations might be missed in these protein-protein interaction studies, the combined unbiased interactomics analyses of USP7 yields a crucial set of high-confidence USP7 partners and substrates.

### Substrate selection by the USP7 TRAF and UBL domains

To assess which domains of USP7 (Fig. 2*A*) are involved in binding partner proteins, we performed a series of IP-MS experiments on WCEs of HEK293T cells transfected with Flag-tagged USP7 deletion constructs. All IPs were performed in three biological replicate experiments, followed by on-bead digestion and tryptic peptide analysis by data-dependent acquisition (DDA)-LFQ-MS. Notably, the catalytic domain (CD) by itself does not bind any of USP7’s interactors (Fig. 2*B* and Supplemental Table S3). In contrast, the combined TRAF and CD domains bind a subset of interactors, including ACTL8, DAXX, DDX24, FAM172A and SCML2. For DNMT1 and GMPS, the UBLs, but not the TRAF domain, are necessary and sufficient for binding to USP7. Finaly, the association of some proteins, such as MAGED4, MAGED2, PJA1, PPIL4 and ncPRC1.1 and ncPRC1.6, involves both the TRAF domain and the UBLs. Note that in the MS analysis peptides were detected over a wide dynamic intensity range and, thus, the resulting protein abundances based on MaxQuant iBAQ values vary by several orders of magnitude (reflected in the shades of blue in the color map).

**Fig. 2.**
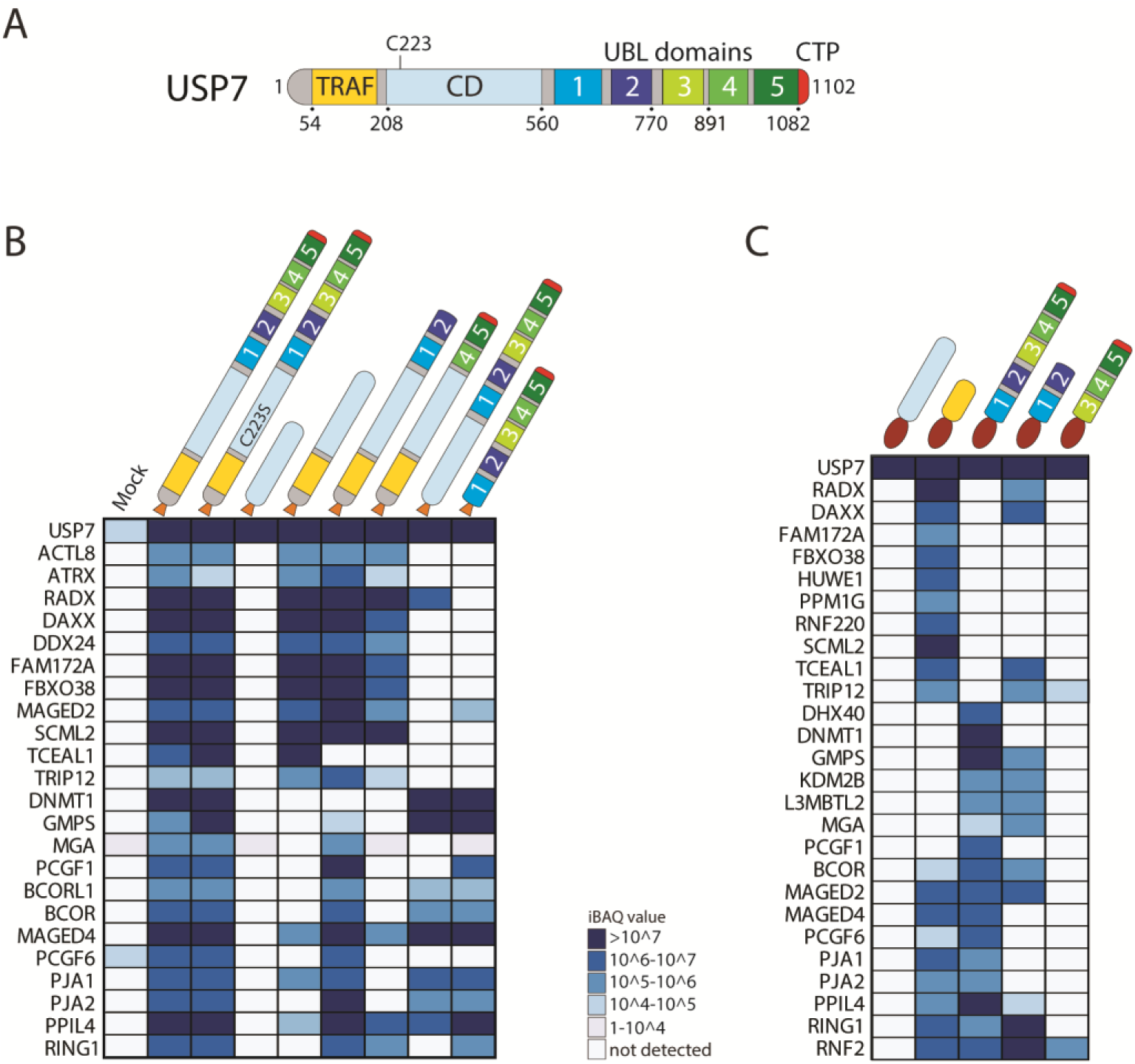
Substrate binding by USP7 TRAF and UBL domains. ***A***, Schematic presentation of the multi-domain organization of USP7. The TRAF domain, catalytic domain (CD), ubiquitin-like (UBL) domains and C-terminal peptide (CTP) are indicated. ***B***, Interaction heatmap of high-confidence USP7 partner protein binding to the indicated Flag-tagged USP7 versions expressed in HEK293T cells, identified by DDA-LFQ-MS analysis of three biological replicate experiments. For the complete data set see Supplemental Table S3. ***C***, Interaction heatmap of high-confidence USP7 partner protein binding to the indicated GST-tagged USP7 domains, following incubation with HEK293T WCEs. Proteins were identified by DDA-LFQ-MS analysis of three biological replicate experiments. For the complete data set see Supplemental Table S4.

As a complementary approach, we compared protein binding to purified gluthatione-S-Transferase (GST)-tagged CD, TRAF domain, UBL1-5 UBL1-2 and UBL3-5 (Fig. 2*C* and Supplemental Table S4). Again, we observed no binding to the CD domain and a clear separation between TRAF and UBL-binding factors. Additionally, a selection of USP7 targets bound both the TRAF domain and the UBLs. Efficient association of PCGF1, BCOR, DHX40, DNMT1 and PPIL4 required all 5 UBLs, whereas binding of MGA, L3MBTL2 and KDM2B mainly relied on UBL1-2. In contrast, UBL4-5 did not appear to play a major role in substrate binding. Taken together, these results show that substrate selection by USP7 is mediated by the TRAF or UBL domains. Particularly UBL1-2 are involved in protein-protein interactions. Its multi-domain arrangement allows USP7 to interact with a variety of targets, and might enable differential regulation of distinct substrates.

### Impact of USP7 inhibition on the ubiquitinome and proteome

To determine the proteome-wide effects of deubiquitylation by USP7, we used the selective inhibitor FT671 for acute inactivation of USP7 (47). HEK293T cells were either mock (7 mM DMSO) treated, or 10 µM FT671 was added for 1 or 6 h. Following trypsin digestion, conjugated ub leaves its C-terminal diglycine remnant on the epsilon amino group of target lysine residues (K-GG). We took advantage of a selective antibody that allows the IP of peptides harboring this K-GG motif, which were next identified by DIA-LFQ-MS. We will refer to the identified K-GGs as ubiquitylation sites (although a small minority of K-GGs reflect modifications by ubiquitin-like proteins NEDD8 or ISG15). In parallel, we determined the impact of USP7 inhibition (USP7i) on the overall proteome by DIA-LFQ-MS. All experiments were performed in three biological replicates per condition. In total, we identified 31,341 K-GG peptides following IP-MS analysis of samples from either mock-treated or cells exposed to FT671. To monitor changes in ubiquitylated peptide abundance, we first filtered the K-GG peptide enriched dataset for peptides with measured LFQ values in all three replicates in at least one experimental condition and then performed a statistical double-sided t-test with Benjamini-Hochberg FDR (0.05) used for truncation (Fig. 3*A*-*B* and Supplemental Table S5-6). We identified 2,267 and 3,941 ubiquitylation sites that increased, and 1,912 and 599 that decreased after treatment with FT671 for 1 or 6 h, respectively (defined as peptides with a LOG2 T-test difference of <-1.00 or >1.00 and p-value <0.05). K-GG peptides that changed significantly are indicated in blue, whereas those derived from high-confidence USP7 interactors are shown in red. Following USP7i, we observed increased ubiquitylation of all high confidence interactors. We note that Steger et al (44), also identified BRWD2/PHIP, BRWD3, FBXO38, L3MBTL2, MGA, PCGF6, RNF220, TCEAL4, TOPORS and TRIM27 as potential USP7 targets that were rapidly ubiquitylated and destabilized following USP7i. As a specific example, MGA-derived K-GG peptides are shown in orange. Already after 1 hr, USP7i leads to a dramatic increase in ubiquitylation of USP7-binding proteins, including virtually all modification sites on MGA. Finally, USP7i did not have a notable impact on the various ubiquitylation sites within ubiquitin itself (Supplemental Fig. S2 and Supplemental Table S5).

**Fig. 3.**
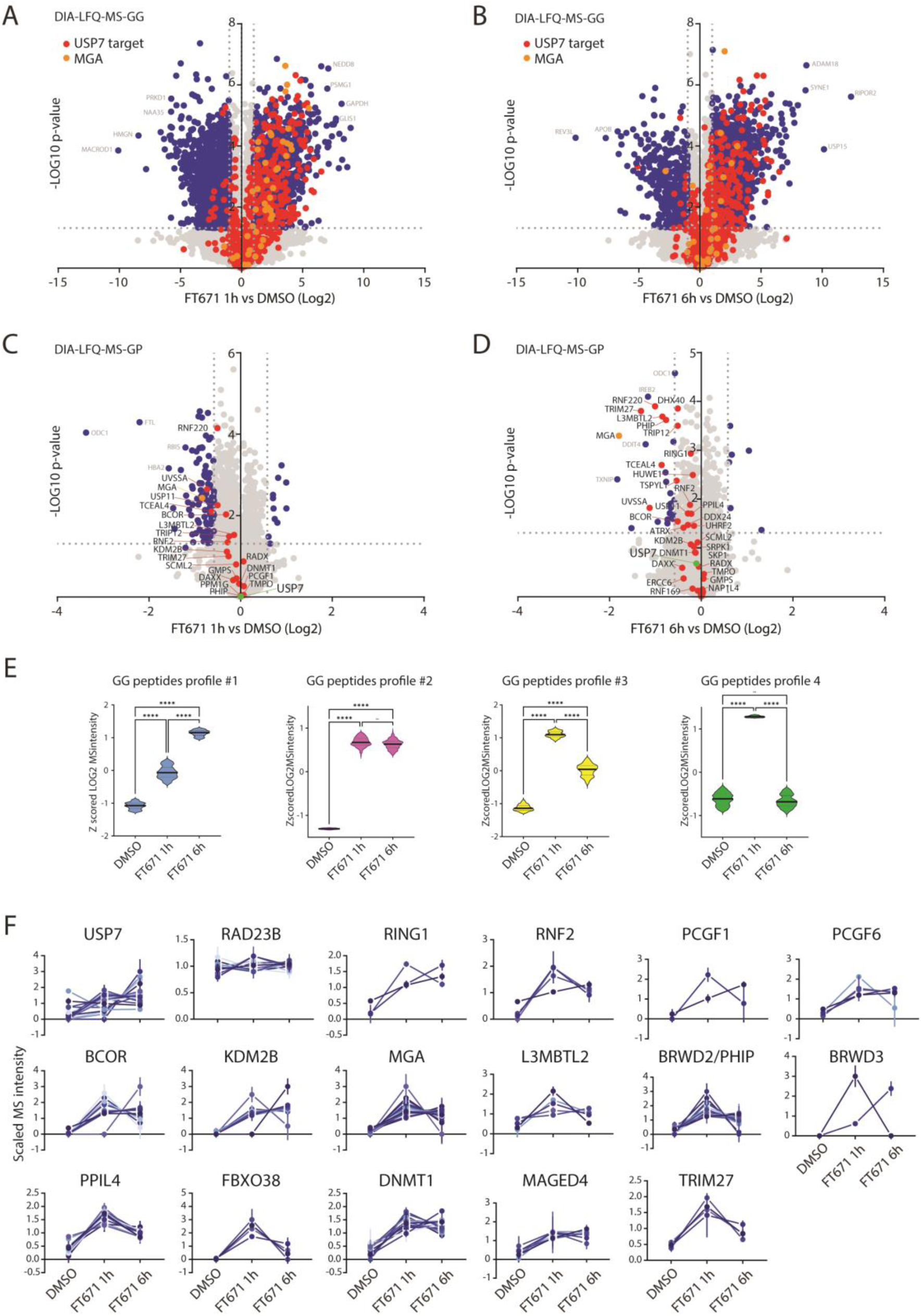
Impact of USP7i on the ubiquitinome and proteome. ***A*-*B***, Volcano plots depicting the ubiquitinome of HEK293T cells after (*A*) 1 or (*B*) 6 h in the presence of 10 µM FT671 (dissolved in DMSO, 7mM final concentration) versus 6 h in the presence of 7 mM DMSO. After enrichment by IP, K-GG peptides were identified by DIA-LFQ-MS in three biological replicate experiments. The K-GG peptide data set was filtered for peptides with measured LFQ values in all three replicates in at least one experimental condition, and then selected by a statistical double-sided t-test with Benjamini-Hochberg FDR (0.05) used for truncation (Fig. 3A-*B* and Supplemental Table S5-6). After 1 hr of USP7i, 2,267 K-GG peptides increased in abundance, whereas 1,912 decreased. Following 6 h of USP7i, 3,941 ubiquitylation sites increased and 599 decreased (defined as peptides with a LOG2 T-test difference of <-1.00 or >1.00 and p-value <0.05). K-GG peptides that changed significantly are indicated in blue, those derived from USP7 partner proteins shown in red and MGA-derived K-GG peptides are shown in orange. ***C*-*D***, Volcano plots depicting the proteome of HEK293T cells after (C) 1 or (D) 6 h in the presence of FT671 versus 6 h in the presence of DMSO. Proteins were identified by DIA-LFQ-MS in three biological replicate experiments. The peptide data set was filtered for peptides with measured LFQ values in all three replicates in at least one experimental condition, and then selected by a statistical double-sided t-test with Benjamini-Hochberg FDR (0.05) used for truncation. Outliers were defined as proteins with a Log2 t test difference of <-1.00 or >1.00 and a p-value of <0.05. Proteins with a significant change in abundance are indicated in blue, high confidence USP7 interacting proteins in red, USP7 in green and MGA in orange. Note that a selection of putative USP7 target proteins were subjected to PRM-MS to obtain higher quantitative accuracy (see Supplemental Fig. S3). ***E***, Violin plots for the top 100 or top 200 peptides with the lowest Pearson distance to one of the four *a priori* defined reference profiles, based on fuzzy C means clustering of Z scored transformed LFQ data. The median is represented by the solid black line, the quartiles by the dashed black lines. ***F***, Ubiquitylation abundance plots of a selection of high-confidence USP7 targets. Intensities of K-GG peptides within a substrate that were identified by DIA-LFQ-MS were normalized for protein abundance determined by either DIA-LFQ-MS or PRM-MS. RAD23B, which is not a substrate of USP7, serves as a control.

While USP7i had a substantial effect on the ubiquitinome within the 1 to 6 h treatment with FT671, the overall impact on the global proteome was small (Fig. 3*C*-*D* and Supplemental Table S5,7). However, the abundance of some USP7 substrates, e.g., BCOR, BRWD2/PHIP, L3MBTL2, MGA, PCGF6, RNF220, TCEAL4, TRIM27 and UVSSA, was reduced significantly. In contrast, USP7i had no appreciable effect on the levels of other partner proteins such as DAXX, DNMT1, GMPS, TMPO and RADX. For proteins with a long half-life, the full effect of USP7i on abundance might well take longer than 6 h (see e.g. references (25, 26, 28). We note also that inclusion in the volcano plot requires the presence of valid LFQ intensity values in all samples, since no data imputation was applied in our workflow. Consequently, proteins such as BRWD3, FBXO38 and TOPORS whose abundances were below the detection limit following USP7i, were not included in the global proteome plots (but are analyzed below).

To further explore the dynamics of protein ubiquitylation in response to USP7i, we performed an unbiased profile analysis to reveal trends in K-GG peptide abundance (Fig. 3*E* and Supplemental Table S8). This analysis is based on fuzzy C means clustering of Z scored transformed data, in which measured K-GG peptide abundance is compared to *a priori* defined reference profiles, based on the lowest Pearson distance. We defined four reference profiles to capture differential ubiquitylation in response to USP7i: 1) peptides with gradually increasing K-GG abundance; 2) high levels of K-GG measured at 1 hr USP7i that persist; 3) strong increase of K-GG at 1 hr USP7i that is reduced at 6 h; 4) strong increase of K-GG at 1 hr USP7i that returns to base level after 6 h of USP7i. This profile analysis revealed that the USP7i has different effects on distinct ubiquitylation sites. To explore this notion in more detail, we next plotted K-GG abundance for each site within a substrate, normalized for protein levels (Fig. 3*F* and Supplemental Table S7). Again, this analysis showed that different ubiquitylation sites (sometimes within the same substrate) responded markedly different to USP7i. Taken together, our proteome-wide analysis of the impact of USP7i on the ubiquitinome and proteome suggests that the effects of USP7 activity are surprisingly substrate and ubiquitylation site-specific.

### Auto-deubiquitylation of USP7

Next, we analyzed the impact of USP7i on the ubiquitylation and abundance of high-confidence substrates in more detail. Although below we focus on deubiquitylation of proteins bound by USP7, we note that our ubiquitinomics analysis also reveals potential additional targets that were missed in the protein-interaction assays. In order to optimize the quantitative accuracy, we used parallel reaction monitoring (PRM)-MS as a complementary method to the proteome-wide DIA-LFQ-MS, to determine the effects of USP7i on protein abundance of the majority of USP7 substrates (Supplemental Fig. S3-4 and Supplemental Table S7-9). Ubiquitylation of USP7 itself increased dramatically following USP7i, suggesting loss of auto-deubiquitylation (Fig. 4*A*-*B*, Supplemental Fig. S3-4 and Supplemental Table S7-9). In Fig. 4*A*, we indicated the 15 ubiquitylation sites detected in USP7 that increased more than 2-fold in the presence of FT671. Notably, many of these ubiquitylation sites within USP7 are located in or near UBL3. The height of the lollipop bars reflects the relative abundance of the corresponding K-GG peptides based on the LFQ values. Note that the K-GG peptide intensity values that were measured vary over multiple orders of magnitude, suggesting large differences in ubiquitylation stoichiometries. USP7i has no substantial effect on the overall abundance of USP7 (Fig. 4*B*, red bars; Supplemental Fig. S3). However, already after 1 h of USP7i there is a massive increase in ubiquitylation of multiple Ks in USP7 (Fig. 4*B*, blue bars; Supplemental Fig. S3-4 and Supplemental Table S7,9). Note that all plotted ubiquitylation levels in the bar plots are normalized for overall protein abundance (calculated based on non-modified tryptic peptide intensities only). The K-GG peptide associated with K1096 of USP7 and located within the CTP is among the peptides with the highest absolute intensity values, suggesting that the ubiquitylation stoichiometry on this residue is relatively high. It will be interesting to explore the possibility that (de)ubiquitylation of K1096 plays a role in activation of the CD by the CTP. In contrast, USP7i did not substantially affect the ubiquitylation of RAD23B, which is not a substrate of USP7, and shown here only as a reference (Fig. 4*B*, Supplemental Fig. S3-4 and Supplemental Table S7,9).

**Fig. 4.**
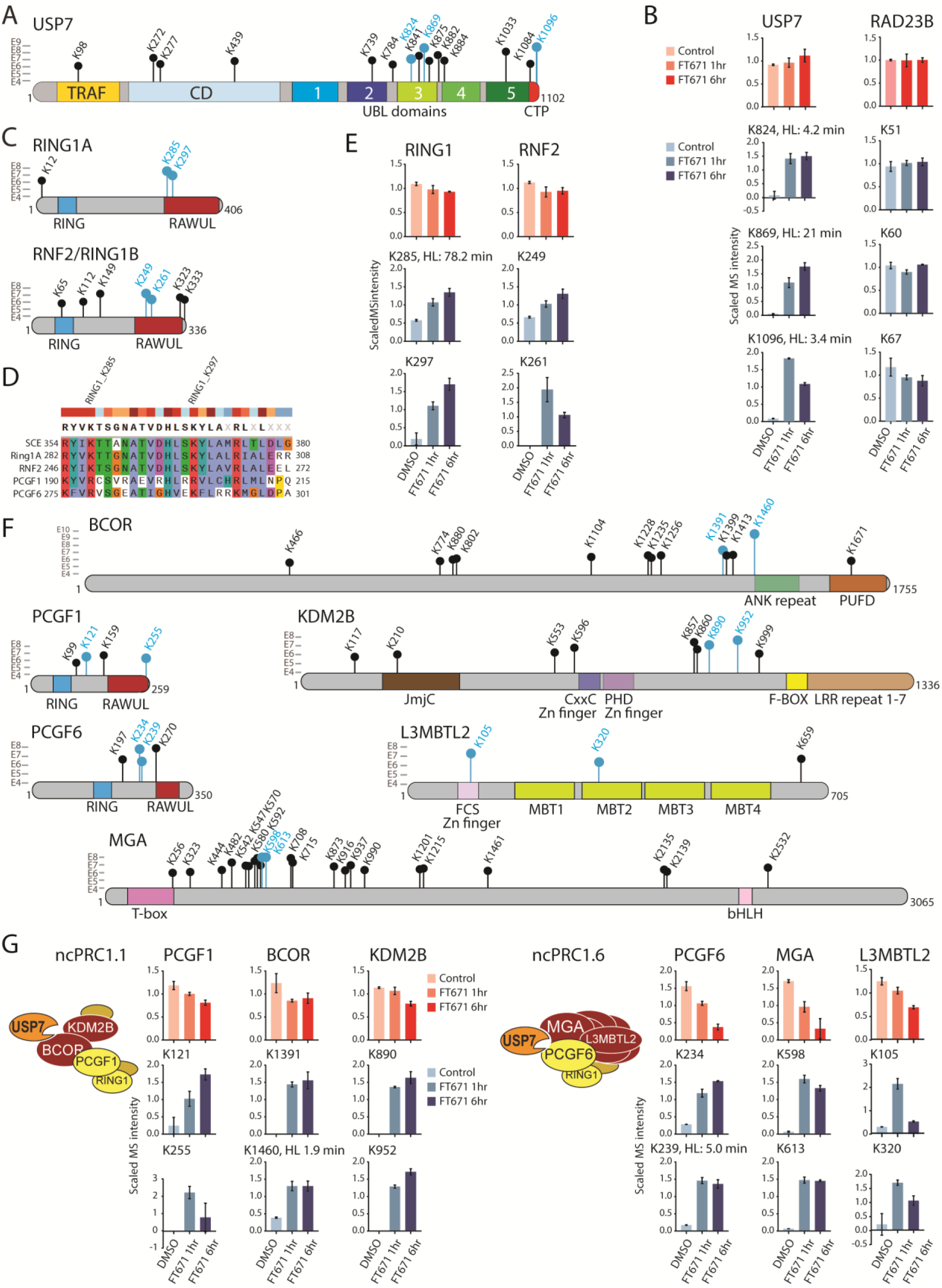
Effect of USP7-dependent deubiquitylation on the Polycomb system. ***A***, Schematic presentation of USP7i-sensitive ubiquitylation sites within USP7. Ubiquitylation sites that change more than two-fold upon USP7i are indicated as lollipops. The height of the lollipop bars reflects the relative abundance of the corresponding K-GG peptides based on the LFQ values, indicated by the Log10 scale bar on the side. Blue lollipops indicate ubiquitylation sites that are represented in the bar-graphs in panel *B*. ***B***, Bar graphs of scaled relative MS intensities representing protein abundances (red) in HEK293T cells cultured in the presence of 7 mM DMSO (6 h), or 10 µM FT671 for 1 or 6 h. Corresponding scaled relative MS intensities representing K-GG peptide abundances that are normalized for overall protein abundance are shown in blue bar graphs. RAD23B, which is not a substrate of USP7, serves as a control. Note that only a small selection of USP7-regulated ubiquitylation sites is shown, for a complete set of K-GG peptides per protein, see Supplemental Fig. S3-4 and Supplemental Table S7,9. Half-lives (HL) of ubiquitylation are based on (48). ***C***, USP7i-sensitive ubiquitylation within RING1A and RNF2/RING1B. ***D***, Alignment of part of the conserved RAWUL domain harboring ubiquitylated Ks in human RNG1,RNF2, PCGF1 and PCGF6 and *Drosophila* Sex Combs Extra (SCE). ***E***, Protein abundance and ubiquitylation of RING1A and RNF2 in response to USP7i. ***F***, USP7i-sensitive ubiquitylation sites within key subunits of ncPRC1.1 (PCGF1, BCOR and KDM2B) and ncPRC1.6 (PCGF6, MGA and L3MBTL2). Ankyrin repeat (ANK repeat); Zinc finger (Zn finger); Leucine rich repeat (LRR); Malignant Brain Tumor (MBT); basic helix-loop-helix (bHLH); PCGF Ub-like fold discriminator (PUFD); RING finger- and WD40-associated ubiquitin-like domain (RAWUL); Jumonji C-domain (JmjC). ***G***, Representative selection of USP7i-sensitive ubiquitylation of ncPRC1.1 and ncPRC1.6. Bar graphs as in *B*. Note that only a small selection of USP7-regulated ubiquitylation sites is shown, for a complete set of K-GG peptides per protein, see Supplemental Fig. S3-4 and Supplemental Table S7,9. Half-life (HL).

Recently, a systems-scale determination of site-selective deubiquitylation rates, following inhibition of the E1 ubiquitin activating step, revealed that the median half-life of global ubiquitylation (in HeLa cells) is ∼12 min (48). However, ubiquitin turnover at individual sites was found to vary by over two orders of magnitude, ranging from less than 1 minute to over 2 h. We used the data set provided by Prus et al. (48) to relate the USP7-regulated ubiquitylation sites to ubiquitin turnover rates. We identified three USP7i-sensitive ubiquitylation sites within USP7 in the data set of Prus et al., which had half-lives ranging from 3.4 to 21 min (Fig. 4B). In conclusion, USP7 undergoes cycles of ubiquitylation and auto-deubiquitylation that are highly site-specific and with different dynamics.

### Impact of deubiquitylation by USP7 on the Polycomb system

USP7 associates with ncPRC1.1 and ncPRC1.6, and is crucial for their integrity and stability (25–28). However, the specifics and kinetics of deubiquitylation of these complexes by USP7 remained undetermined. Therefore, we examined the impact of USP7i on the ubiquitylation and abundance of ncPRC1.1 and ncPRC1.6. Note that all plotted ubiquitylation levels in the bar plots are normalized for protein abundance. Although RING1A and RING1B are only associated with USP7 when they reside within either ncPRC1.1 or ncPRC1.6 (26), USP7i has a notable effect on their ubiquitylation status (Fig. 4*C* and Supplemental Fig. S3-4 and Supplemental Table S7,9). Particularly, the level of ubiquitylation of two conserved Ks within the variant ubiquitin-fold RAWUL domain of RING1A and RING1B is induced following USP7i _(_Fig. 4*D*-*E*, Supplemental Fig. S3-4 and Supplemental Table S7,9). As the RAWUL domain is a crucial protein-protein interaction domain, its (de)ubiquitylation might modulate the association of partner proteins. Ubiquitylation of signature subunits of ncPRC1.1 (PCGF1, BCOR and KDM2B) and ncPRC1.6 (PCGF6, MGA and L3MBTL2) is strongly upregulated during USP7i (Fig. 4*F*-*G*, Supplemental Fig. S3-4 and Supplemental Table S7,9), consistent with the notion that they are all direct target sites of USP7. ncPRC1.1 is destabilized following long-term USP7i (>1 day) and in cells lacking USP7 (26, 28). However, following 6 h in the presence of FT671, the overall levels of ncPRC1.1 subunits PCGF1, BCOR and KDM2B were only modestly reduced (Fig. 4*G*, Supplemental Fig. S3-4 and Supplemental Table S7,9). In contrast, the overall protein abundance of key ncPRC1.6 subunits MGA and PCGF6 is strongly reduced following USP7i. Thus, ncPRC1.6 is more immediately destabilized by USP7i than ncPRC1.1, while the protein levels of other PRC1s and PRC2 are not affected by the loss of USP7 activity (Supplemental Fig. S3-4 and Supplemental Table S7,9; see also (26, 28)).

In conclusion, our analysis revealed a range of substrate-specific effects of USP7 on its Polycomb targets. USP7 controls the dosage of ncPRC1.1 and ncPRC1.6, albeit with markedly different kinetics. The integrity of ncPRC1.6 is more acutely dependent on deubiquitylation by USP7 than that of ncPRC1.1. It will also be important to evaluate the effect of ubiquitylation on Polycomb complex activity when protein stability itself is not affected. E.g., the (de)ubiquitylation of key functional domains of USP7 or RING1A/B are likely to play a regulatory role. The high-resolution map of USP7-regulated ubiquitylation sites we presented here, provides an essential resource and starting point for such functional studies.

### Substrate-specific dynamics of deubiquitylation by USP7

We extended our analysis of the impact of USP7 activity on the ubiquitylation status and stability of additional high-confidence partner proteins (Fig. 5, Supplemental Fig. S3-4 and Supplemental Table S7-9). BRWD2/PHIP was previously identified as a USP7 target that requires USP7 for its stability (25). Indeed, we found that USP7i leads to increased ubiquitylation and reduced levels of BRWD2/PHIP, albeit with relatively slow kinetics (Fig. 5*A*-*B*, Supplemental Fig. S3-4 and Supplemental Table S7,9). In the present study, we also identified BRWD1 and BRWD3 as a novel substrate of USP7. Following USP7i, BRWD3 is ubiquitylated and its protein levels reduced rapidly. The effects of USP7i on BRWD1 stability were intermediate between BRWD2/PHIP and BRWD3 (Supplemental Fig. S3). Thus, similar to PCGF1 and PCGF6, the homologous BRWD1, BRWD2/PHIP and BRWD3 are stabilized by USP7, but with substantially different half-lives. Although PPIL4 protein abundance is hardly affected within the 6 h time-frame of the experiment, there is a rapid increase in ubiquitylation following USP7i. We also note that the half-life of K212 ubiquitylation was measured at 4.5 min, whereas K321 ubiquitylation has a half-life of over 120 min (48). DNMT1 protein abundance is not affected by USP7i, but ubiquitylation of a large number of Ks increased substantially. Again, the turnover rate of different USP7-regulated sites within DNMT1 can vary by about an order of magnitude. For other substrates, such as FBXO38, MAGED4 and TRIM27, deubiquitylation by USP7 is crucial for protein stability (Fig. 5*A*-*B*, Supplemental Fig. S3-4 and Supplemental Table S7,9). However, the ubiquitylation patterns following USP7i can vary substantially between targets or even between different sites within the same substrate (Fig. 4-5, Supplemental Fig. S3-4 and Supplemental Table S7,9). After one hour in the presence of FT671, we could no longer detect overall TOPORS levels, indicating that the stability of this protein is highly dependent upon deubiquitylation by USP7 (Fig. 5*A*-*B*, Supplemental Fig. S3-4 and Supplemental Table S7,9).

**Fig. 5.**
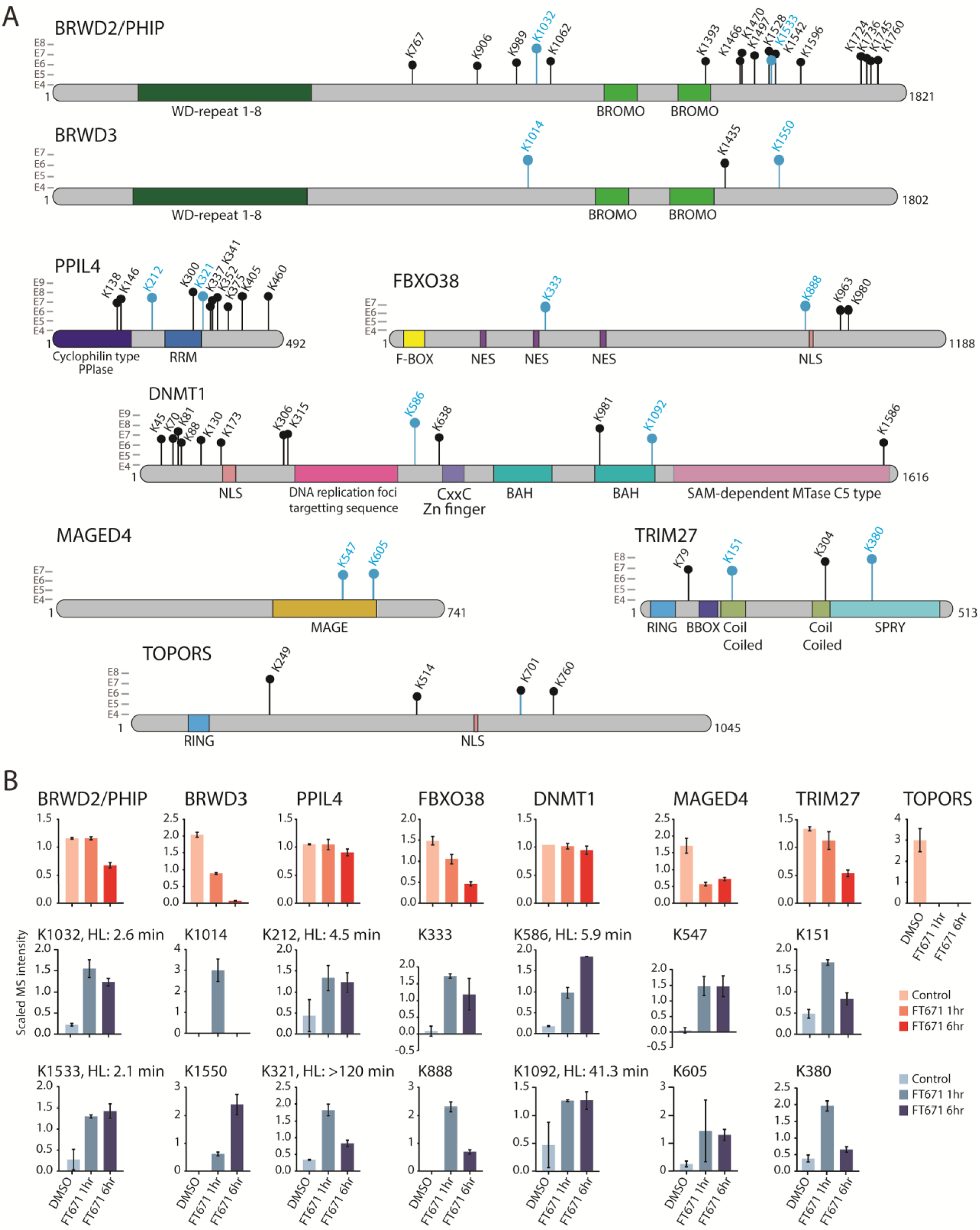
Substrate-specific effects of deubiquitylation by USP7. ***A***, Schematic presentation of USP7i-sensitive ubiquitylation sites within a selection of substrates. Annotation as described for Fig. 4A. RNA recognition motif (RRM); nuclear export signal (NES); Nuclear localization signal (NLS); bromo-adjacent homology (BAH); S-adenosylmethionine (SAM); methyltransferase (MTase); peptidyl-prolyl cis-trans isomerase (PPIase). ***B***, Bar graphs representing scaled relative MS intensities representing protein abundances (red) and scaled relative MS intensities representing K-GG peptide abundances (blue) that are normalized for overall protein abundance, in HEK293T cells cultured in the presence of 7 mM DMSO (6 h), or 10 µM FT671 for 1 or 6 h. For a complete set of K-GG peptides per protein, see Supplemental Fig. S3-4 and Supplemental Table S7,9. Half-lives (HL) of ubiquitylation are based on (48).

In summary, we provided a detailed proteome-wide map of sites directly or indirectly deubiquitylated by USP7 and its consequences for protein abundances of interaction partners and target/substrates. Notably, our analysis revealed that USP7’s activity profile is highly substrate-specific. Even different USP7-regulated ubiquitylation sites within a protein can display differential kinetic responses to USP7i. Finally, increased ubiquitylation stoichiometry in response to USP7i can have different consequences for the stability of different substrates.

## DISCUSSION

In the present study we took advantage of recent advances in MS-based ubiquitinomics to perform an unbiased survey of USP7 targets. First, we combined the results of our IP-DIA-LFQ-MS analysis of endogenous USP7 with earlier IP-MS analyses of epitope-tagged USP7 (from our and other labs) to obtain a consensus high-confidence interactome. The USP7 network comprises a variety of multi-protein hubs, dedicated to distinct nuclear functions. Second, we found that both the TRAF domain and UBLs contribute to substrate selectivity of USP7. This multi-domain arrangement might enable differential regulation of separate substrates. Third, we determined the impact of rapid chemical inhibition of USP7 on the ubiquitinome, yielding a high-resolution proteome-wide map of USP7 target sites. Fourth, we found that (de)ubiquitylation kinetics of USP7 vary between substrates and is not an immutable enzymatic property. Fifth, increased ubiquitylation after USP7i destabilizes some substrates but not others. Because some USP7-regulated ubiquitylation sites are within key functional domains, they might regulate substrate activity, independent of its stability. In summary, we provided a detailed map of the dynamic impact of USP7 on the ubiquitinome and found that USP7 enzymatic activity and effects are highly substrate-specific.

We defined a consensus, high-confidence set of USP7 substrates, involved in chromatin dynamics, Polycomb gene repression, RNA processing, DNA-damage response and transcription control (Fig. 1*G*). Several USP7 targets form subcomplexes dedicated to specific functions. E.g., TCEAL1/USP11 is involved in transcriptional elongation (49). A separate hub is formed by the E3 ubiquitin ligase PJA1, associates with MAGED1/2/4, the histone chaperone NAP1 and the DUB USP9X. Accumulating evidence suggests that PJA1 acts as a suppressor of a broad range of protein-aggregation associated with neurodegenerative disease (50). Moreover, Praja1 might connect to the Polycomb system through ubiquitylation-mediated regulation of PRC2 (51). We also discovered a physical association between the DNA repair factor RADX and the E3 ubiquitin ligase TRIM27 (see also (26)). USP7 binds and regulates the H2AK119 ubiquitin ligase complexes ncPRC1.1 and ncPRC1.6, but not the related ncPRC1.3/5 or cPRC1.2/4.

Another striking feature is USP7’s association with multiple E3 ubiquitin ligases (thirteen), ubiquitin adaptor proteins (seven) and one other DUB, USP11. Several of these E3s, e.g. TOPORS, PCGF6, PCGF1 and TRIM27, require deubiquitylation by USP7 for their stability. Conversely, associated E3s ubiquitylate USP7, which is counteracted by auto-deubiquitylation. While ubiquitylation of USP7 does not affect its abundance, it is likely to modulate its activity or set of interactions, because ubiquitylation occurs within key functional domains (Fig. 4*A*-*B*). The regulatory implications of the frequent pairing of the DUB USP7 with various E3 ubiquitin ligases remain to be explored further (9, 52). We also note that some USP7 partner proteins might directly modulate its activity, similar to GMP-synthase (12, 20, 33, 34). Finally, in our aim to select a set of high-confidence USP7 targets we might well have missed transient, condition-dependent or cell-type-specific interactions.

USP7 is a remarkably pleiotropic DUB. A main research focus has been on USP7i as a means to activate the p53 tumor suppressor pathway. However, unbiased studies of the USP7 network (25, 26, 28, 31, 32, 44), including the present study, revealed additional tumor suppressors and oncoproteins that are targeted by USP7 (Fig. 6). MGA, as part of ncPRC1.6, opposes the tumor-driving activity of MYC (53, 54). Likewise, BCOR is a tumor suppressor implicated in acute myeloid leukemia, T-cell leukemias, sarcomas and neuronal tumors (55). Depending on the type of cancer, HUWE1 and TRIP12 have been described as tumor suppressors but also as cancer drivers (56, 57). TOPORS is a potential tumor suppressor, whose expression is often decreased in cancer cells (58). However, targeting TOPORS augments the cytotoxicity of several chemotherapeutics in cancer cells (58–61). Because TOPORS abundance is highly dependent on USP7 activity (Fig. 5*B*), USP7i might similarly increase the efficacy of anti-cancer therapeutics. Notably, TOPORS, HUWE1 and TRIP12 have also been linked to the p53 tumor suppressor pathway (56–58). PHIP/BRWD2 is an oncogene that is mutated in acute myeloid leukemia, particularly in people of African descent (62, 63). Finally, mutations in *USP7* itself have been implicated in pediatric T-cell acute lymphoblastic leukemia (64). In conclusion, USP7 acts as a rheostat for multiple oncogenic and tumor suppressor pathways. A point of concern is that potential benefits of USP7i for one pathway might be accompanied by undesired effects in another one. Therefore, a deeper understanding of the USP7 regulatory network will be essential to guide the potential therapeutic use of USP7 inhibitors.

**Fig. 6.**
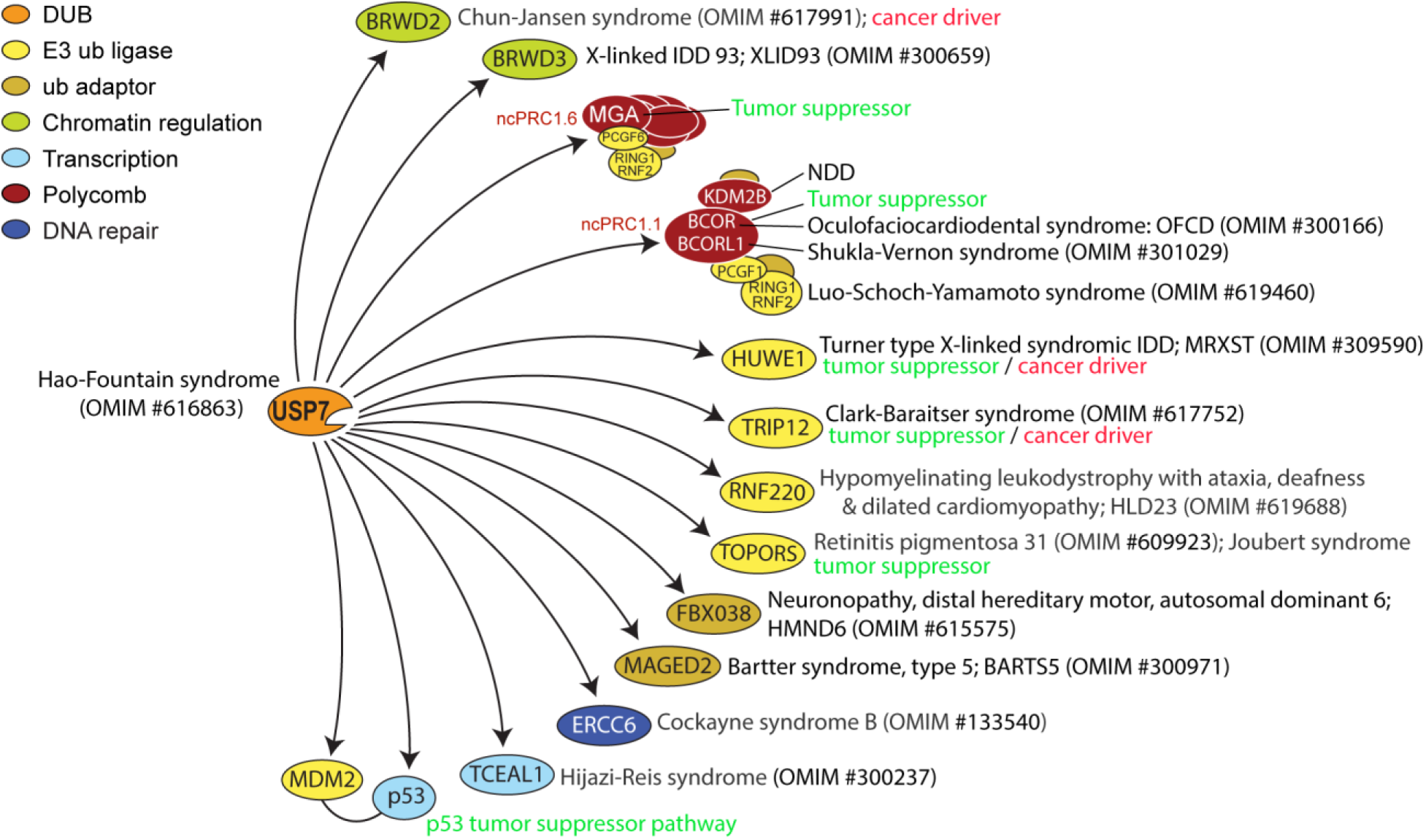
USP7 connects various neurodevelopmental syndromes. Schematic summary of the high number of USP7 targets that are associated with various neurodevelopmental disorders (NDDs). Molecular function, associated syndromes and OMIM numbers are indicated. For details see the main text.

Haploinsufficiency of USP7 causes Hao-Fountain syndrome, a NDD characterized by global developmental delay, impaired intellectual and speech development, behavioral anomalies, including autism spectrum disorder, eye anomalies, seizures and dysmorphic features (14–16). Recently, we found that ncPRC1.1 mediates a large portion of USP7-dependent gene regulation that underpins neuronal differentiation (28). In contrast, ncPRC1.6 does not appear to play a substantial role in neuronal differentiation, reflecting functional diversification between distinct ncPRC1s (28). Indeed, mutations in multiple ncPRC1.1 subunits, but not in ncPRC1.6, cause a variety of neurodevelopmental disorders (53, 55, 65–71) (Fig. 6). A striking number of additional USP7 substrates, including PHIP/BRWD2 (72), BRWD3 (73), HUWE1 (74), TRIP12 (75), RNF220 (76), TOPORS (77), FBXO38 (78, 79), MAGED2 (80, 81), ERCC6 (82) and TCEAL1 (83, 84), are associated with distinct NDDs (Fig. 6). Thus, Hao-Fountain syndrome is likely to be the result of the cumulative effects of reduced functioning of multiple USP7 substrates. Our identification of the core USP7 regulome provides the basis for a systematic dissection of the molecular basis of Hao-Fountain syndrome and related NDDs.

Our results provided unexpected insights into the kinetics and consequences of deubiquitylation by USP7. The time-dependent changes in ubiquitylation following USP7i was remarkably different between different sites (sometimes within the same substrate). Extrapolating from a systems-scale determination of ubiquitylation turnover rates (48), we estimated that ubiquitin half-lives at USP7-regulated sites can vary by well over an order of magnitude (from ∼1 to >120 min). Ubiquitylation status is the resultant of ligation and deubiquitylation. In addition to substrate site-specific context, the close pairing of USP7 with different E3 ubiquitin ligases is likely to play an important role in ubiquitylation dynamics. The abundance of only a subset of substrates is affected by deubiquitylation by USP7, sometimes with substantial differences in protein half-lives. Finally, through targeting of key functional domains, USP7-regulated ubiquitylation could modulate substrate functionality, without affecting its abundance. Besides the substrates of USP7 that were selected and described based on our extensive interaction network analysis, the unbiased ubiquitinome analysis revealed additional potential, yet uncharacterized, USP7 targets (Fig. 3). The K-GG peptides with dynamics that are consistent with the defined ubiquitylation regulation profile plots are among the most prominent responders of USP7 inhibition and are therefore expected to be directly or indirectly regulated by USP7. Although here we have focused on substrates that are readily identified through direct binding to USP7, our comprehensive ubiquitinome analysis is a resource for follow-up research that may reveal additional *bona fide* targets of USP7. In conclusion, our detailed mapping of USP7 ubiquitinome and proteome dynamics, provides a crucial starting point to explore how it connects diverse tumor suppressor, cancer and neurodevelopmental pathways.

## Supporting information

Wolf2025-supplement

## ACKNOWLEDGEMENTS

We are grateful to our lab members for helpful discussions. We thank Ben Tilly and Yaser Atlasi for critical reading of the manuscript.

## Author contributions

Conceptualization: J.vdK., J.W.-vdM., J.A.A.D., A.S. and C.P.V. initiated and designed the study. J.W.-vdM., J.vdK. and A.S. conducted the cell and biochemical experiments, K.B., D.H.W.D., W.A.S.D. and J.A.A.D. performed mass spectrometry and proteomics data analysis. J.W.-vdM., J.vdK., J.A.A.D. and C.P.V. co-wrote the manuscript. All authors discussed results and commented on the manuscript.

## DATA AVAILABILITY

The mass spectrometry proteomics data have been deposited to the ProteomeXchange Consortium via the PRIDE partner repository with the data set identifier PXD071773. All other data needed to evaluate the conclusions in this paper are present in the paper and Supplemental Materials.

## SUPPLEMENTAL DATA

This article contains supplemental data.

## REFERENCES

1. Oh, E., Akopian, D., and Rape, M. (2018) Principles of Ubiquitin-Dependent Signaling. Annu. Rev. Cell Dev. Biol. 34, 137–162

2. Clague, M. J., Heride, C., and Urbé, S. (2015) The demographics of the ubiquitin system. Trends Cell Biol. 25, 417–426

3. Zheng, N., and Shabek, N. (2017) Ubiquitin Ligases: Structure, Function, and Regulation. Annu. Rev. Biochem. 86, 129–157

4. Popovic, D., Vucic, D., and Dikic, I. (2014) Ubiquitination in disease pathogenesis and treatment. Nat. Med. 20, 1242–1253

5. Hershko, A., and Ciechanover, A. (1998) The ubiquitin system. Annu. Rev. Biochem. 67, 425–479

6. Song, L., and Luo, Z. Q. (2019) Post-translational regulation of ubiquitin signaling. J. Cell Biol. 218, 1776–1786

7. Clague, M. J., Urbé, S., and Komander, D. (2019) Breaking the chains: deubiquitylating enzyme specificity begets function. Nat. Rev. Mol. Cell Biol. 20, 338–352

8. Basar, M. A., Beck, D. B., and Werner, A. (2021) Deubiquitylases in developmental ubiquitin signaling and congenital diseases. Cell Death Differ. 28, 538–556

9. Kim, R. Q., and Sixma, T. K. (2017) Regulation of USP7: A High Incidence of E3 Complexes. J. Mol. Biol. 429, 3395–3408

10. Rawat, R., Starczynowski, D. T., and Ntziachristos, P. (2019) Nuclear deubiquitination in the spotlight: the multifaceted nature of USP7 biology in disease. Curr. Opin. Cell Biol. 58, 85–94

11. Pozhidaeva, A., and Bezsonova, I. (2019) USP7: Structure, substrate specificity, and inhibition. DNA Repair (Amst). 76, 30–39

12. Van der Knaap, J. A., Kumar, B. R. P., Moshkin, Y. M., Langenberg, K., Krijgsveld, J., Heck, et al. (2005) GMP synthetase stimulates histone H2B deubiquitylation by the epigenetic silencer USP7. Mol. Cell 17, 695–707

13. Kon, N., Kobayashi, Y., Li, M., Brooks, C. L., Ludwig, T., and Gu, W. (2010) Inactivation of HAUSP in vivo modulates p53 function. Oncogene 29, 1270

14. Hao, Y. H., Fountain, M. D., Fon Tacer, K., Xia, F., Bi, W., Kang, S. H. L., et al. (2015) USP7 Acts as a Molecular Rheostat to Promote WASH-Dependent Endosomal Protein Recycling and Is Mutated in a Human Neurodevelopmental Disorder. Mol. Cell 59, 956–969

15. Fountain, M. D., Oleson, D. S., Rech, M. E., Segebrecht, L., Hunter, J. V., McCarthy, J. M., et al. (2019) Pathogenic variants in USP7 cause a neurodevelopmental disorder with speech delays, altered behavior, and neurologic anomalies. Genet. Med. 21, 1797–1807

16. Wimmer, M. C., Brennenstuhl, H., Hirsch, S., Dötsch, L., Unser, S., Caro, P., et al. (2024) Hao-Fountain syndrome: 32 novel patients reveal new insights into the clinical spectrum. Clin. Genet. 105, 499–509

17. Korchak, E. J., Sharafi, M., Maisonet, I. J., Salazar-Chaparro, A., Semenova, I. V, Khan, H., et al. (2025) Functional spectrum of USP7 pathogenic variants in Hao–Fountain syndrome: Insights into the enzyme’s activity, stability, and allosteric modulation. Proc. Natl. Acad. Sci. 122, e2510252122

18. Schauer, N. J., Liu, X., Magin, R. S., Doherty, L. M., Chan, W. C., Ficarro, S. B., et al. (2020) Selective USP7 inhibition elicits cancer cell killing through a p53-dependent mechanism. Sci. Rep. 10, 5324

19. Brooks, C. L., Li, M., Hu, M., Shi, Y., and Gu, W. (2007) The p53--Mdm2—HAUSP complex is involved in p53 stabilization by HAUSP. Oncogene 26, 7262–7266

20. Reddy, B. A., van der Knaap, J. A., Bot, A. G. M., Mohd-Sarip, A., Dekkers, D. H. W., Timmermans, M. A., et al. (2014) Nucleotide Biosynthetic Enzyme GMP Synthase Is a TRIM21-Controlled Relay of p53 Stabilization. Mol. Cell 53, 458–470

21. Kon, N., Zhong, J., Kobayashi, Y., Li, M., Szabolcs, M., Ludwig, T., et al. (2011) Roles of HAUSP-mediated p53 regulation in central nervous system development. Cell Death Differ. 18, 1366

22. Gao, Z., Zhang, J., Bonasio, R., Strino, F., Sawai, A., Parisi, F., et al. (2012) PCGF Homologs, CBX Proteins, and RYBP Define Functionally Distinct PRC1 Family Complexes. Mol. Cell 45, 344–356

23. Kang, H., Cabrera, J. R., Zee, B. M., Kang, H. A., Jobe, J. M., Hegarty, M. B., et al. (2022) Variant Polycomb complexes in Drosophila consistent with ancient functional diversity. Sci. Adv. 8, eadd0103

24. Lecona, E., Narendra, V., and Reinberg, D. (2015) USP7 cooperates with SCML2 to regulate the activity of PRC1. Mol. Cell. Biol. 35, 1157–1168

25. Nie, L., Wang, C., Liu, X., Teng, H., Li, S., Huang, M., et al. (2022) USP7 substrates identified by proteomics analysis reveal the specificity of USP7. Genes Dev. 36, 1016–1030

26. Sijm, A., Atlasi, Y., van der Knaap, J. A., Wolf van der Meer, J., Chalkley, G. E., Bezstarosti, K., et al. (2022) USP7 regulates the ncPRC1 Polycomb axis to stimulate genomic H2AK119ub1 deposition uncoupled from H3K27me3. Sci. Adv. 8, eabq7598

27. Maat, H., Atsma, T. J., Hogeling, S. M., Rodríguez López, A., Jaques, J., Olthuis, M., et al. (2021) The USP7-TRIM27 axis mediates non-canonical PRC1.1 function and is a druggable target in leukemia. iScience 24, 102435

28. Wolf van der Meer, J., Larue, A., van der Knaap, J. A., Chalkley, G. E., Sijm, A., Beikmohammadi, L., et al. (2025) Hao-Fountain syndrome protein USP7 controls neuronal differentiation via BCOR-ncPRC1.1. Genes Dev. 39, 401–422

29. Schwertman, P., Lagarou, A., Dekkers, D. H. W., Raams, A., van der Hoek, A. C., Laffeber, C., et al. (2012) UV-sensitive syndrome protein UVSSA recruits USP7 to regulate transcription-coupled repair. Nat. Genet. 44, 598

30. Zhang, X., Horibata, K., Saijo, M., Ishigami, C., Ukai, A., Kanno, S., et al. (2012) Mutations in UVSSA cause UV-sensitive syndrome and destabilize ERCC6 in transcription-coupled DNA repair. Nat. Genet. 44, 593

31. Georges, A., Coyaud, E., Marcon, E., Greenblatt, J., Raught, B., and Frappier, L. (2019) USP7 Regulates Cytokinesis through FBXO38 and KIF20B. Sci. Rep. 9, 2724

32. Georges, A., Marcon, E., Greenblatt, J., and Frappier, L. (2018) Identification and Characterization of USP7 Targets in Cancer Cells. Sci. Rep. 8, 15833

33. van der Knaap, J. A., Kozhevnikova, E., Langenberg, K., Moshkin, Y. M., and Verrijzer, C. P. (2010) Biosynthetic Enzyme GMP Synthetase Cooperates with Ubiquitin-Specific Protease 7 in Transcriptional Regulation of Ecdysteroid Target Genes. Mol. Cell. Biol. 30, 736–744

34. Faesen, A. C., Dirac, A. M. G., Shanmugham, A., Ovaa, H., Perrakis, A., and Sixma, T. K. (2011) Mechanism of USP7/HAUSP activation by its C-terminal ubiquitin-like domain and allosteric regulation by GMP-synthetase. Mol. Cell 44, 147–159

35. Park, H.-B., and Baek, K.-H. (2023) Current and future directions of USP7 interactome in cancer study. Biochim. Biophys. Acta - Rev. Cancer 1878, 188992

36. Sheng, Y., Saridakis, V., Sarkari, F., Duan, S., Wu, T., Arrowsmith, C. H., et al. (2006) Molecular recognition of p53 and MDM2 by USP7/HAUSP. Nat. Struct. Mol. Biol. 13, 285–291

37. Bojagora, A., and Saridakis, V. (2020) USP7 manipulation by viral proteins. Virus Res. 286, 198076

38. Rougé, L., Bainbridge, T. W., Kwok, M., Tong, R., Di Lello, P., Wertz, I. E., et al. (2016) Molecular Understanding of USP7 Substrate Recognition and C-Terminal Activation. Structure 24, 1335–1345

39. Kim, R. Q., Geurink, P. P., Mulder, M. P. C., Fish, A., Ekkebus, R., El Oualid, F., et al. (2019) Kinetic analysis of multistep USP7 mechanism shows critical role for target protein in activity. Nat. Commun. 10, 1–16

40. Jaen Maisonet, I., Sharafi, M., Korchak, E. J., Salazar-Chaparro, A., Bratt, A. S., Parikh, T., et al. (2025) Small-molecule allosteric activator of ubiquitin-specific protease 7 (USP7). Proc. Natl. Acad. Sci. U. S. A. 122, e2510496122

41. Korchak, E. J., Geddes-Buehre, D. H., and Bezsonova, I. (2025) The C-terminal Tail of Ubiquitin-Specific Protease 7 Facilitates Ubiquitin Release and Ensures Efficient Catalytic Turnover. J. Mol. Biol. 437, 169298

42. Pfoh, R., Lacdao, I. K., Georges, A. A., Capar, A., Zheng, H., Frappier, L., et al. (2015) Crystal Structure of USP7 Ubiquitin-like Domains with an ICP0 Peptide Reveals a Novel Mechanism Used by Viral and Cellular Proteins to Target USP7. PLOS Pathog. 11, 1–23

43. Bushman, J. W., Donovan, K. A., Schauer, N. J., Liu, X., Hu, W., Varca, A. C., et al. (2021) Proteomics-Based Identification of DUB Substrates Using Selective Inhibitors. Cell Chem. Biol. 28, 78–87.e3

44. Steger, M., Demichev, V., Backman, M., Ohmayer, U., Ihmor, P., Müller, S., et al. (2021) Time-resolved in vivo ubiquitinome profiling by DIA-MS reveals USP7 targets on a proteome-wide scale. Nat. Commun. 12, 5399

45. Chalkley, G. E., and Verrijzer, C. P. (2004) Immuno-depletion and purification strategies to study chromatin-remodeling factors in vitro. Methods Enzymol. 377, 421–442

46. Tyanova, S., Temu, T., Sinitcyn, P., Carlson, A., Hein, M. Y., Geiger, T., et al. (2016) The Perseus computational platform for comprehensive analysis of (prote)omics data. Nat. Methods 13, 731–740

47. Turnbull, A. P., Ioannidis, S., Krajewski, W. W., Pinto-Fernandez, A., Heride, C., Martin, A. C. L., et al. (2017) Molecular basis of USP7 inhibition by selective small-molecule inhibitors. Nature 550, 481–486

48. Prus, G., Satpathy, S., Weinert, B. T., Narita, T., and Choudhary, C. (2024) Global, site-resolved analysis of ubiquitylation occupancy and turnover rate reveals systems properties. Cell 187, 2875–2892.e21

49. Dehmer, M., Trunk, K., Gallant, P., Fleischhauer, D., Müller, M., Herold, S., et al. (2025) The USP11/TCEAL1 complex promotes transcription elongation to sustain oncogenic gene expression in neuroblastoma. Genes Dev. 39, 751–769

50. Watabe, K., Niida-Kawaguchi, M., Tada, M., Kato, Y., Murata, M., Tanji, K., et al. (2022) Praja1 RING-finger E3 ubiquitin ligase is a common suppressor of neurodegenerative disease-associated protein aggregation. Neuropathology 42, 488–504

51. Zoabi, M., Sadeh, R., de Bie, P., Marquez, V. E., and Ciechanover, A. (2011) PRAJA1 is a ubiquitin ligase for the polycomb repressive complex 2 proteins. Biochem. Biophys. Res. Commun. 408, 393–398

52. Bolhuis, D. L., Emanuele, M. J., and Brown, N. G. (2024) Friend or foe? Reciprocal regulation between E3 ubiquitin ligases and deubiquitinases. Biochem. Soc. Trans. 52, 241–267

53. Mathsyaraja, H., Catchpole, J., Freie, B., Eastwood, E., Babaeva, E., Geuenich, M., et al. (2021) Loss of MGA repression mediated by an atypical polycomb complex promotes tumor progression and invasiveness. Elife 10, e64212

54. Llabata, P., Mitsuishi, Y., Choi, P. S., Cai, D., Francis, J. M., Torres-Diz, M., et al. (2020) Multi-Omics Analysis Identifies MGA as a Negative Regulator of the MYC Pathway in Lung Adenocarcinoma. Mol. Cancer Res. 18, 574–584

55. Astolfi, A., Fiore, M., Melchionda, F., Indio, V., Bertuccio, S. N., and Pession, A. (2019) BCOR involvement in cancer. Epigenomics 11, 835–855

56. Brunet, M., Vargas, C., Larrieu, D., Torrisani, J., and Dufresne, M. (2020) E3 Ubiquitin Ligase TRIP12: Regulation, Structure, and Physiopathological Functions. Int. J. Mol. Sci. 21, 8515

57. Kao, S.-H., Wu, H.-T., and Wu, K.-J. (2018) Ubiquitination by HUWE1 in tumorigenesis and beyond. J. Biomed. Sci. 25, 67

58. Ji, L., Huo, X., Zhang, Y., Yan, Z., Wang, Q., and Wen, B. (2020) TOPORS, a tumor suppressor protein, contributes to the maintenance of higher-order chromatin architecture. Biochim. Biophys. acta. Gene Regul. Mech. 1863, 194518

59. Truong, P., Shen, S., Joshi, S., Islam, M. I., Zhong, L., Raftery, M. J., et al. (2024) TOPORS E3 ligase mediates resistance to hypomethylating agent cytotoxicity in acute myeloid leukemia cells. Nat. Commun. 15, 7360

60. Jaffray, E. G., Tatham, M. H., Mojsa, B., Plechanovová, A., Rojas-Fernandez, A., Liu, J. C. Y., et al. (2025) PML mutants from arsenic-resistant patients reveal SUMO1-TOPORS and SUMO2/3-RNF4 degradation pathways. J. Cell Biol. 224, e202407133

61. Kaito, S., Aoyama, K., Oshima, M., Tsuchiya, A., Miyota, M., Yamashita, M., et al. (2024) Inhibition of TOPORS ubiquitin ligase augments the efficacy of DNA hypomethylating agents through DNMT1 stabilization. Nat. Commun. 15, 7359

62. Wu, X., and Levine, R. L. (2025) A new link in the leukemia genetic puzzle. Genes Dev. 39, 1129–1131

63. Pawar, A. S., Somers, P., Alex, A., Grana, J., Feist, V. K., George, S. S., et al. (2025) Leukemia mutated proteins PHF6 and PHIP form a chromatin complex that represses acute myeloid leukemia stemness. Genes Dev. 39, 1219–1240

64. Shaw, T. I., Dong, L., Tian, L., Qian, C., Liu, Y., Ju, B., et al. (2021) Integrative network analysis reveals USP7 haploinsufficiency inhibits E-protein activity in pediatric T-lineage acute lymphoblastic leukemia (T-ALL). Sci. Rep. 11, 5154

65. Ryan, C. W., Peirent, E. R., Regan, S. L., Guxholli, A., and Bielas, S. L. (2024) H2A monoubiquitination: insights from human genetics and animal models. Hum. Genet. 143, 511–527

66. Tamburri, S., Conway, E., and Pasini, D. (2022) Polycomb-dependent histone H2A ubiquitination links developmental disorders with cancer. Trends Genet. 38, 333–352

67. Pierce, S. B., Stewart, M. D., Gulsuner, S., Walsh, T., Dhall, A., McClellan, J. M., et al. (2018) De novo mutation in RING1 with epigenetic effects on neurodevelopment. Proc. Natl. Acad. Sci. U. S. A. 115, 1558–1563

68. Shukla, A., Girisha, K. M., Somashekar, P. H., Nampoothiri, S., McClellan, R., and Vernon, H. J. (2019) Variants in the transcriptional corepressor BCORL1 are associated with an X-linked disorder of intellectual disability, dysmorphic features, and behavioral abnormalities. Am. J. Med. Genet. A 179, 870–874

69. van Jaarsveld, R. H., Reilly, J., Cornips, M.-C., Hadders, M. A., Agolini, E., Ahimaz, P., et al. (2023) Delineation of a KDM2B-related neurodevelopmental disorder and its associated DNA methylation signature. Genet. Med. 25, 49–62

70. Hamline, M. Y., Corcoran, C. M., Wamstad, J. A., Miletich, I., Feng, J., Lohr, J. L., et al. (2020) OFCD syndrome and extraembryonic defects are revealed by conditional mutation of the Polycomb-group repressive complex 1.1 (PRC1.1) gene BCOR. Dev. Biol. 468, 110–132

71. Ragge, N., Isidor, B., Bitoun, P., Odent, S., Giurgea, I., Cogné, B., et al. (2019) Expanding the phenotype of the X-linked BCOR microphthalmia syndromes. Hum. Genet. 138, 1051–1069

72. Morgan, M. A. J., Popova, I. K., Vaidya, A., Burg, J. M., Marunde, M. R., Rendleman, E. J., et al. (2021) A trivalent nucleosome interaction by PHIP/BRWD2 is disrupted in neurodevelopmental disorders and cancer. Genes Dev. 35, 1642–1656

73. Field, M., Tarpey, P. S., Smith, R., Edkins, S., O’Meara, S., Stevens, C., et al. (2007) Mutations in the BRWD3 gene cause X-linked mental retardation associated with macrocephaly. Am. J. Hum. Genet. 81, 367–374

74. De Falco, A., Minale, E. M. P., Meossi, C., Pagano, S., Trovato, R., Agolini, E., et al. (2025) Exploring the Clinical Spectrum of HUWE1-Related Neurodevelopmental Disorder: Five New Patients and Literature Review. Am. J. Med. Genet. A 197, e63959

75. Aerden, M., Denommé-Pichon, A.-S., Bonneau, D., Bruel, A.-L., Delanne, J., Gérard, B., et al. (2023) The neurodevelopmental and facial phenotype in individuals with a TRIP12 variant. Eur. J. Hum. Genet. 31, 461–468

76. Ma, P., and Mao, B. (2022) The many faces of the E3 ubiquitin ligase, RNF220, in neural development and beyond. Dev. Growth Differ. 64, 98–105

77. Strong, A., Qu, H.-Q., Cullina, S., McManus, M. L., Zackai, E. H., Glessner, J., et al. (2023) TOPORS as a novel causal gene for Joubert syndrome. Am. J. Med. Genet. A 191, 2156–2163

78. Sumner, C. J., d’Ydewalle, C., Wooley, J., Fawcett, K. A., Hernandez, D., Gardiner, A. R., et al. (2013) A dominant mutation in FBXO38 causes distal spinal muscular atrophy with calf predominance. Am. J. Hum. Genet. 93, 976–983

79. Pegat, A., Chanson, J.-B., Lozeron, P., Joubert, B., Bani-Sadr, A., Quadrio, I., et al. (2024) Identification of rare variants in the FBXO38 gene of patients with chronic inflammatory demyelinating polyradiculoneuropathy. J. Neuroimmunol. 392, 578381

80. Yan, X., Hu, Y., Zhang, X., Gao, X., Zhao, Y., Peng, H., et al. (2024) Identification of a novel intronic mutation of MAGED2 gene in a Chinese family with antenatal Bartter syndrome. BMC Med. Genomics 17, 23

81. Buffet, A., Filser, M., Bruel, A., Dard, R., Quibel, T., Dubucs, C., et al. (2025) X-linked transient antenatal Bartter syndrome related to MAGED2 gene: Enriching the phenotypic description and pathophysiologic investigation. Genet. Med. Off. J. Am. Coll. Med. Genet. 27, 101217

82. Wang, Y., Chakravarty, P., Ranes, M., Kelly, G., Brooks, P. J., Neilan, E., et al. (2014) Dysregulation of gene expression as a cause of Cockayne syndrome neurological disease. Proc. Natl. Acad. Sci. 111, 14454–14459

83. Albuainain, F., Shi, Y., Lor-Zade, S., Hüffmeier, U., Pauly, M., Reis, A., et al. (2024) Confirmation and expansion of the phenotype of the TCEAL1-related neurodevelopmental disorder. Eur. J. Hum. Genet. 32, 350–356

84. Hijazi, H., Reis, L. M., Pehlivan, D., Bernstein, J. A., Muriello, M., Syverson, E., et al. (2022) TCEAL1 loss-of-function results in an X-linked dominant neurodevelopmental syndrome and drives the neurological disease trait in Xq22.2 deletions. Am. J. Hum. Genet. 109, 2270–2282

