## Supplementary material for "Proteome-wide ubiquitinome profiling reveals substrate-specific dynamics within the USP7 network": Wolf2025-supplement

**Description of Supplemental Tables**

**Supplemental Table S1:** Materials.

**Supplemental Table S2:** MaxQuant search output protein identifications and iBAQ values for all IP-MS experiments.

**Supplemental Table S3:** MaxQuant search output protein identifications and iBAQ values for FLAG-USP7 deletion mutant IP-MS experiments.

**Supplemental Table S4:** MaxQuant search output protein identifications and iBAQ values for GST-USP7 domain AP-MS experiments.

**Supplemental Table S5:** DIA-NN search output data of global proteome (GP) and K-ε-GG *peptide* (GG) proteomics experiments.

**Supplemental Table S6:** Volcano plots data file of GP and GG experiments.

**Supplemental Table S7:** Input data for all DIA-NN output based GP and GG bar graphs.

**Supplemental Table S8:** Profile plots data file of GG categories.

**Supplemental Table S9:** Skyline PRM output data for a selection of target proteins and H2A.

### Supplemental Figures

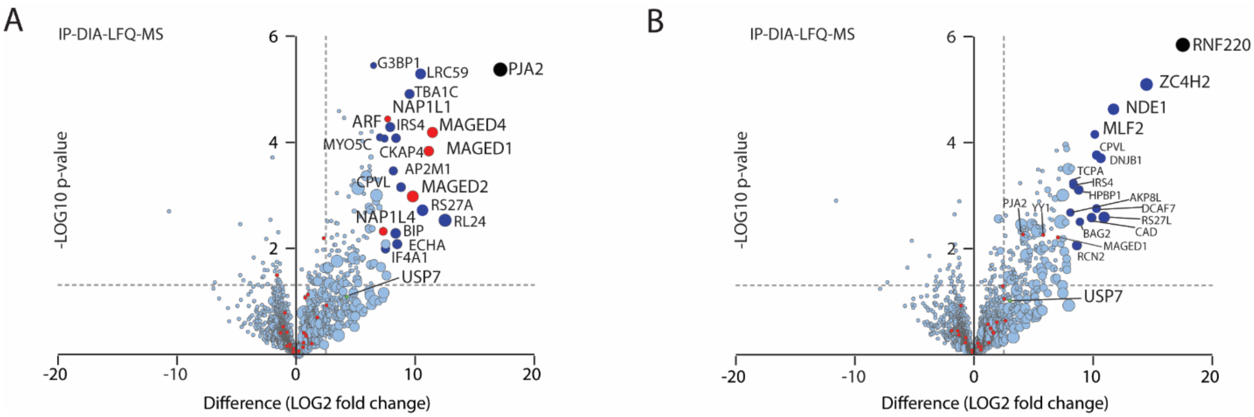


**Fig. S1. PRAJA2 and RNF220 interactome.**

**A-B**, IP-MS analysis of Flag-PJA2 (A) and Flag-RNF220 (B) expressed in HEK293T cells. Volcano plot representing a statistical double-sided t-test based on LFQ quantitative MS data of protein enrichment in IPs from cells expressing a Flag-tagged bait plotted versus IPs from cells that do not. The size of the data spheres reflects the relative abundance of the protein in the IP analysis, based on its mass spectrometry LFQ intensity. The dashed grey lines indicate arbitrary thresholds (i.e., p value = 0.05, LOG2 t test difference = 2 or enrichment over control >4-fold). The analysis is based on three independent biological replicate experiments. High-confidence USP7-interacting proteins are shown in red. The bait is indicated in black. For the complete data set see Supplemental Table S2.

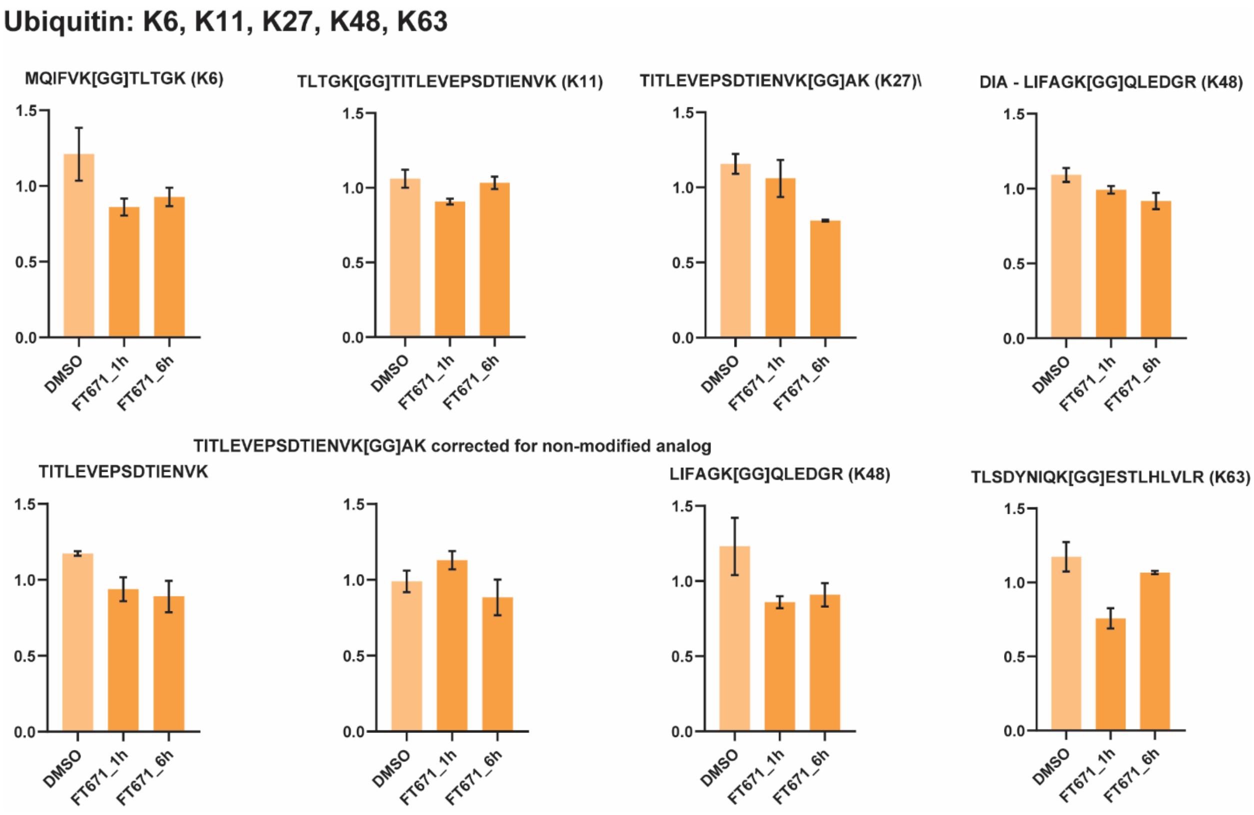


**Fig. S2. Impact of USP7i on ubiquitylation sites within ubiquitin.**
Bar graphs representing scaled relative MS intensities representing K-GG peptide abundances, in HEK293T cells cultured in the presence of 7 mM DMSO (6 h), or 10 µM FT671 for 1 or 6 h, respectively.

**Fig. S3.**

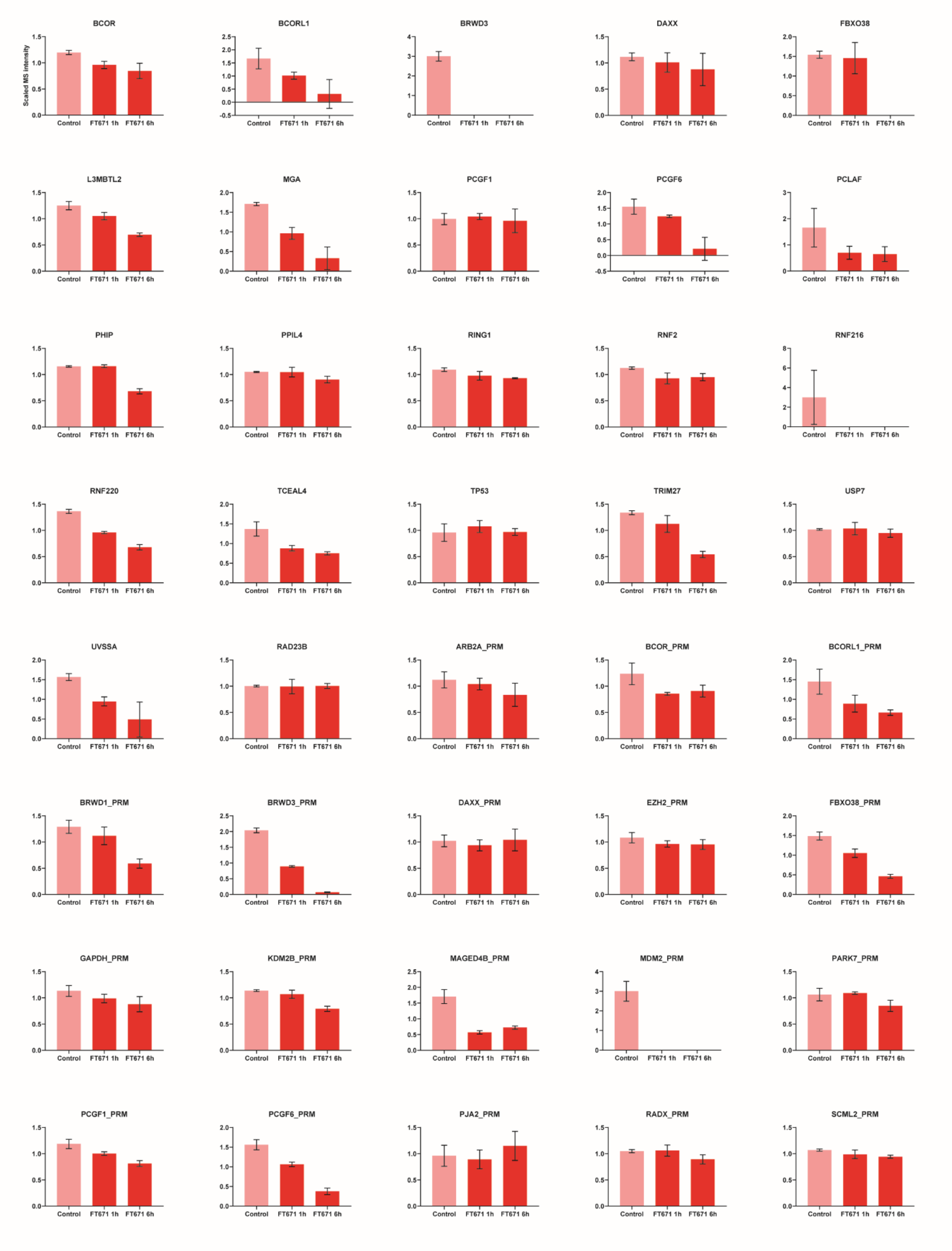
BCOR
BCORL1
BRWD3
DAXX
FBXO38
L3MBTL2
MGA
PCGF1
PCGF6
PCLAF
PHIP
PPIL4
RING1
RNF2
RNF216
RNF220
TCEAL4
TP53
TRIM27
USP7
UVSSA
RAD23B
ARB2A_PRM
BCOR_PRM
BCORL1_PRM
BRWD1_PRM
BRWD3_PRM
DAXX_PRM
EZH2_PRM
FBXO38_PRM
GAPDH_PRM
KDM2B_PRM
MAGED4B_PRM
MDM2_PRM
PARK7_PRM
PCGF1_PRM
PCGF6_PRM
PJA2_PRM
RADX_PRM
SCML2_PRM
Scaled MS intensity
Control
FT671 1h
FT671 6h

**Fig. S3.**

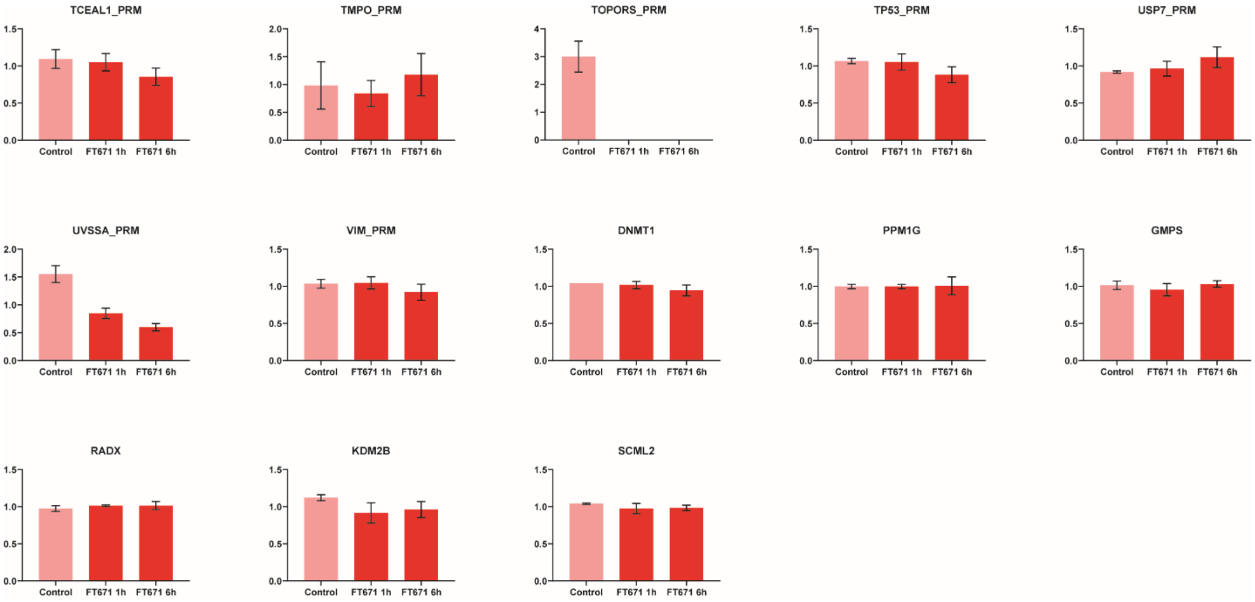


**Fig. S3. Impact of USP7i on the abundance of high-confidence substrates.**
Bar graphs representing scaled relative MS intensities representing protein abundances determined by DIA-LFQ-MS and PRM in HEK293T cells cultured in the presence of 7 mM DMSO (6 h), or 10 µM FT671 for 1 or 6 h. See Supplemental Table S7 for input data for all DIA-NN and Supplemental Table S9 for Skyline PRM output data.

**Fig. S4.**

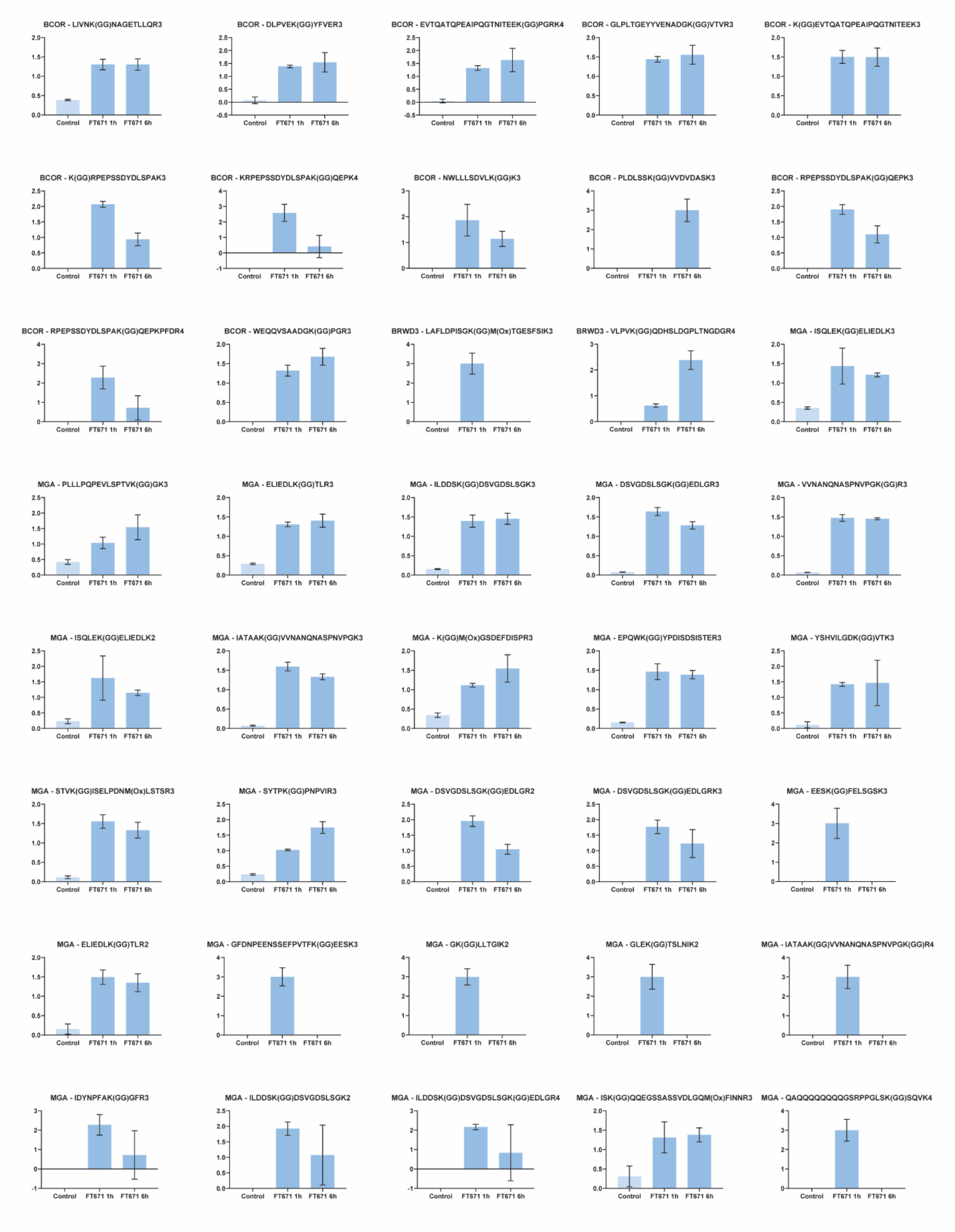
BCOR - LIVNK(GG)NAGETLLQR3
BCOR - DLPVEK(GG)YFVER3
BCOR - EVTQATQPEAIPQGTNITEEK(GG)PGRK4
BCOR - GLPLTGEYYVENADGK(GG)VTVR3
BCOR - K(GG)EVTQATQPEAIPQGTNITEEK3
BCOR - K(GG)RPEPSSDYDLSPAK3
BCOR - KRPEPSSDYDLSPAK(GG)QEPK4
BCOR - NWLLLSDVLK(GG)K3
BCOR - PLDLSSK(GG)VVDVDASK3
BCOR - RPEPSSDYDLSPAK(GG)QEPK3
BCOR - RPEPSSDYDLSPAK(GG)QEPKPFDR4
BCOR - WEQQVSAADGK(GG)PGR3
BRWD3 - LAFLDPISGK(GG)M(Ox)TGESFSIK3
BRWD3 - VLPVK(GG)QDHSLDGPLTNGDGR4
MGA - ISQLEK(GG)ELIEDLK3
MGA - PLLLPQPEVLSPTVK(GG)GK3
MGA - ELIEDLK(GG)TLR3
MGA - ILDDSK(GG)DSVGDSLSGK3
MGA - DSVGDSLSGK(GG)EDLGR3
MGA - VVNANQNASPNVPGK(GG)R3
MGA - ISQLEK(GG)ELIEDLK2
MGA - IATAAK(GG)VVNANQNASPNVPGK3
MGA - K(GG)M(Ox)GSDEFDISPR3
MGA - EPQWK(GG)YPDISDSISTER3
MGA - YSHVILGDK(GG)VTK3
MGA - STVK(GG)ISELPDNM(Ox)LSTSR3
MGA - SYTPK(GG)PNPVIR3
MGA - DSVGDSLSGK(GG)EDLGR2
MGA - DSVGDSLSGK(GG)EDLGRK3
MGA - EESK(GG)FELSGSK3
MGA - ELIEDLK(GG)TLR2
MGA - GFDNPEENSSEFPVTFK(GG)EESK3
MGA - GK(GG)LLTGIK2
MGA - GLEK(GG)TSLNIK2
MGA - IATAAK(GG)VVNANQNASPNVPGK(GG)R4
MGA - IDYNPFAK(GG)GFR3
MGA - ILDDSK(GG)DSVGDSLSGK2
MGA - ILDDSK(GG)DSVGDSLSGK(GG)EDLGR4
MGA - ISK(GG)QQEGSSASSVDLGQM(Ox)FINNR3
MGA - QAQQQQQQQQGSRPPGLSK(GG)SQVK4
Control
FT671 1h
FT671 6h

Fig. S4.

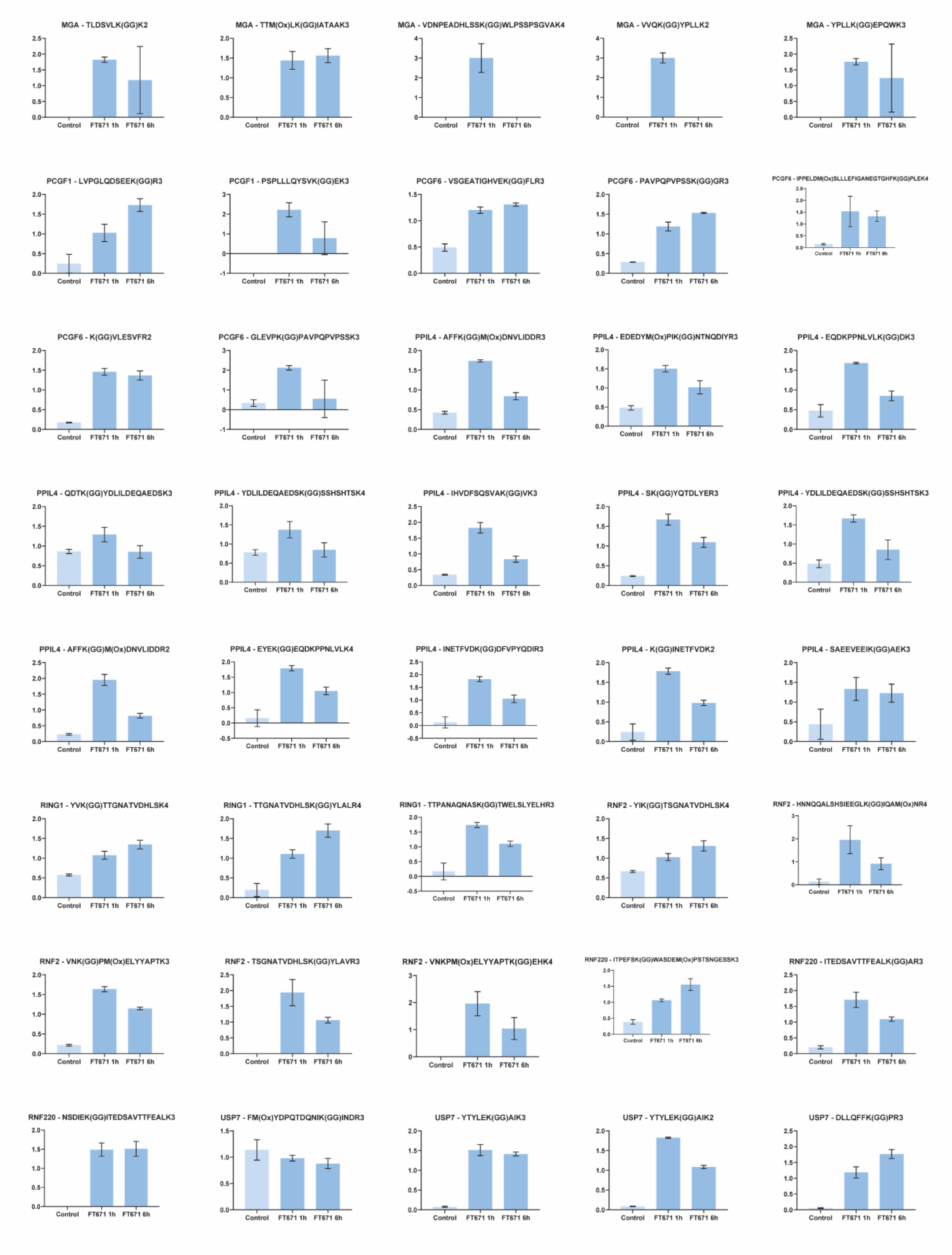
MGA - TLDSVLK(GG)K2
MGA - TTM(Ox)LK(GG)IATAAK3
MGA - VDNPEADHLSSK(GG)WLPSSPSGVAK4
MGA - VVQK(GG)YPLLK2
MGA - YPLLK(GG)EPQWK3
PCGF1 - LVPGLQDSEEK(GG)R3
PCGF1 - PSPLLLQYSVK(GG)EK3
PCGF6 - VSGEATIGHVEK(GG)FLR3
PCGF6 - PAVPQPVPSSK(GG)GR3
PCGF6 - IPPELDM(Ox)SLLLEFIGANEGTGHFK(GG)PLEK4
PCGF6 - K(GG)VLESVFR2
PCGF6 - GLEVPK(GG)PAVPQPVPSSK3
PPIL4 - AFFK(GG)M(Ox)DNVLIDDR3
PPIL4 - EDEDYM(Ox)PIK(GG)NTNQDIYR3
PPIL4 - EQDKPPNLVLK(GG)DK3
PPIL4 - QDTK(GG)YDLILDEQAEDSK3
PPIL4 - YDLILDEQAEDSK(GG)SSHSHTSK4
PPIL4 - IHVDFSQSVAK(GG)VK3
PPIL4 - SK(GG)YQTDLYER3
PPIL4 - YDLILDEQAEDSK(GG)SSHSHTSK3
PPIL4 - AFFK(GG)M(Ox)DNVLIDDR2
PPIL4 - EYEK(GG)EQDKPPNLVLK4
PPIL4 - INETFVDK(GG)DFVPYQDIR3
PPIL4 - K(GG)INETFVDK2
PPIL4 - SAEEVEEIK(GG)AEK3
RING1 - YVK(GG)TTGNATVDHLSK4
RING1 - TTGNATVDHLSK(GG)YLALR4
RING1 - TTPANAQNASK(GG)TWELSLYELHR3
RNF2 - YIK(GG)TSGNATVDHLSK4
RNF2 - HNNQQALSHSIEEGLK(GG)IQAM(Ox)NR4
RNF2 - VNK(GG)PM(Ox)ELYYAPTK3
RNF2 - TSGNATVDHLSK(GG)YLAVR3
RNF2 - VNKPM(Ox)ELYYAPTK(GG)EHK4
RNF220 - ITPEFSK(GG)WASDEM(Ox)PSTSNGESSK3
RNF220 - ITEDSAVTTFEALK(GG)AR3
RNF220 - NSDIEK(GG)ITEDSAVTTFEALK3
USP7 - FM(Ox)YDPQTDQNIK(GG)INDR3
USP7 - YTYLEK(GG)AIK3
USP7 - YTYLEK(GG)AIK2
USP7 - DLLQFFK(GG)PR3
Control
FT671 1h
FT671 6h

**Fig. S4.**

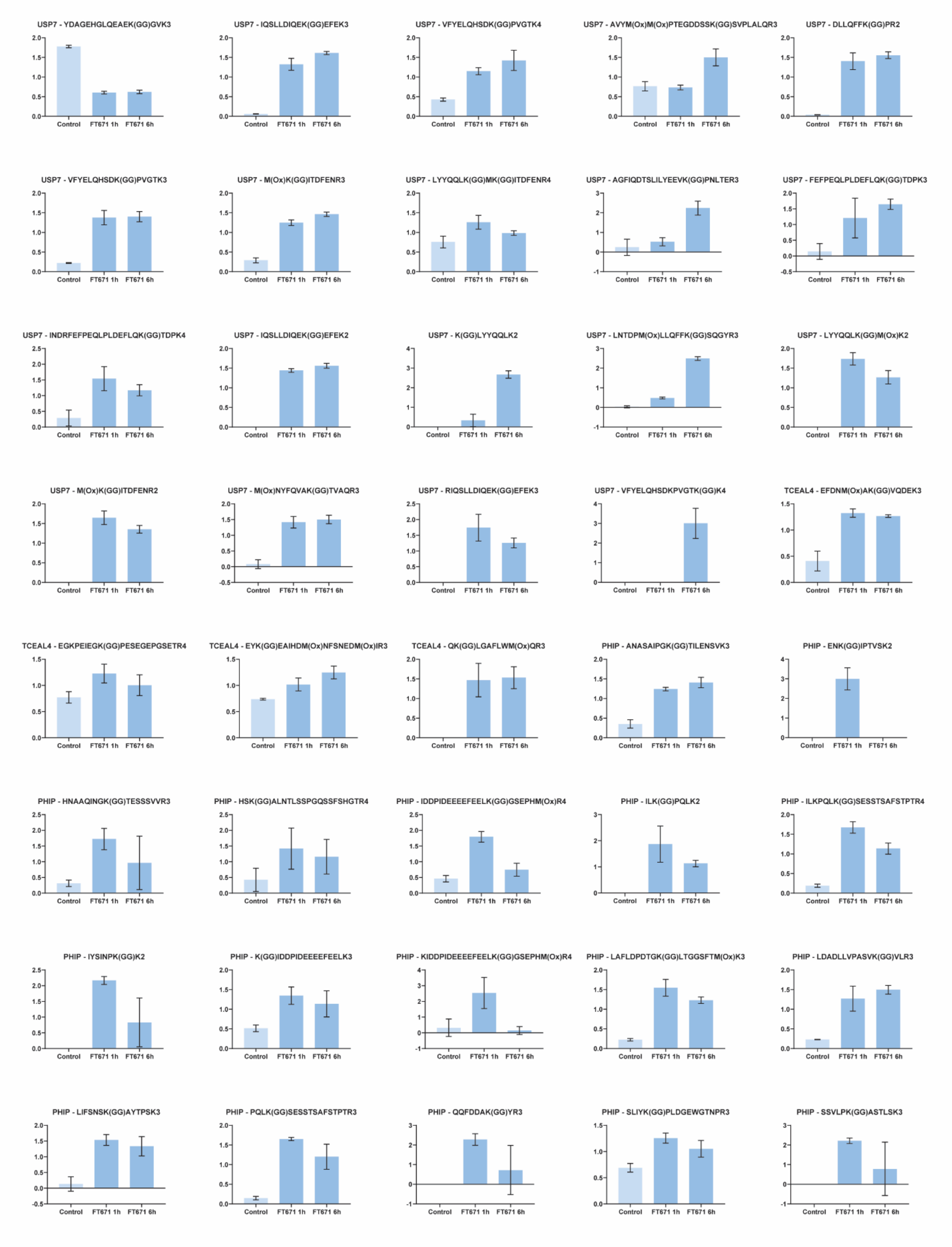
USP7 - YDAGEHGLQEAEK(GG)GVK3
USP7 - IQSLLDIQEK(GG)EFEK3
USP7 - VFYELQHSDK(GG)PVGTK4
USP7 - AVYM(Ox)M(Ox)PTEGDDSSK(GG)SVPLALQR3
USP7 - DLLQFFK(GG)PR2
USP7 - VFYELQHSDK(GG)PVGTK3
USP7 - M(Ox)K(GG)ITDFENR3
USP7 - LYYQQLK(GG)MK(GG)ITDFENR4
USP7 - AGFIQDTSLILYEEVK(GG)PNLTER3
USP7 - FEFPEQLPLDEFLQK(GG)TDPK3
USP7 - INDRFEFPEQLPLDEFLQK(GG)TDPK4
USP7 - IQSLLDIQEK(GG)EFEK2
USP7 - K(GG)LYYQQLK2
USP7 - LNTDPM(Ox)LLQFFK(GG)SQGYR3
USP7 - LYYQQLK(GG)M(Ox)K2
USP7 - M(Ox)K(GG)ITDFENR2
USP7 - M(Ox)NYFQVAK(GG)TVAQR3
USP7 - RIQSLLDIQEK(GG)EFEK3
USP7 - VFYELQHSDKPVGTK(GG)K4
TCEAL4 - EFDNM(Ox)AK(GG)VQDEK3
TCEAL4 - EGKPEIEGK(GG)PESEGEPGSETR4
TCEAL4 - EYK(GG)EAIHDM(Ox)NFSNEDM(Ox)IR3
TCEAL4 - QK(GG)LGAFLWM(Ox)QR3
PHIP - ANASAIPGK(GG)TILENSVK3
PHIP - ENK(GG)IPTVSK2
PHIP - HNAAQINGK(GG)TESSSVVR3
PHIP - HSK(GG)ALNTLSSPGQSSFSHGTR4
PHIP - IDDPIDEEEEFEELK(GG)GSEPHM(Ox)R4
PHIP - ILK(GG)PQLK2
PHIP - ILKPQLK(GG)SESSTSAFSTPTR4
PHIP - IYSINPK(GG)K2
PHIP - K(GG)IDDPIDEEEEFEELK3
PHIP - KIDDPIDEEEEFEELK(GG)GSEPHM(Ox)R4
PHIP - LAFLDPDTGK(GG)LTGGSFTM(Ox)K3
PHIP - LDADLLVPASVK(GG)VLR3
PHIP - LIFSNSK(GG)AYTPSK3
PHIP - PQLK(GG)SESSTSAFSTPTR3
PHIP - QQFDDAK(GG)YR3
PHIP - SLIYK(GG)PLDGEWGTNPR3
PHIP - SSVLPK(GG)ASTLSK3
Control
FT671 1h
FT671 6h

**Fig. S4.**

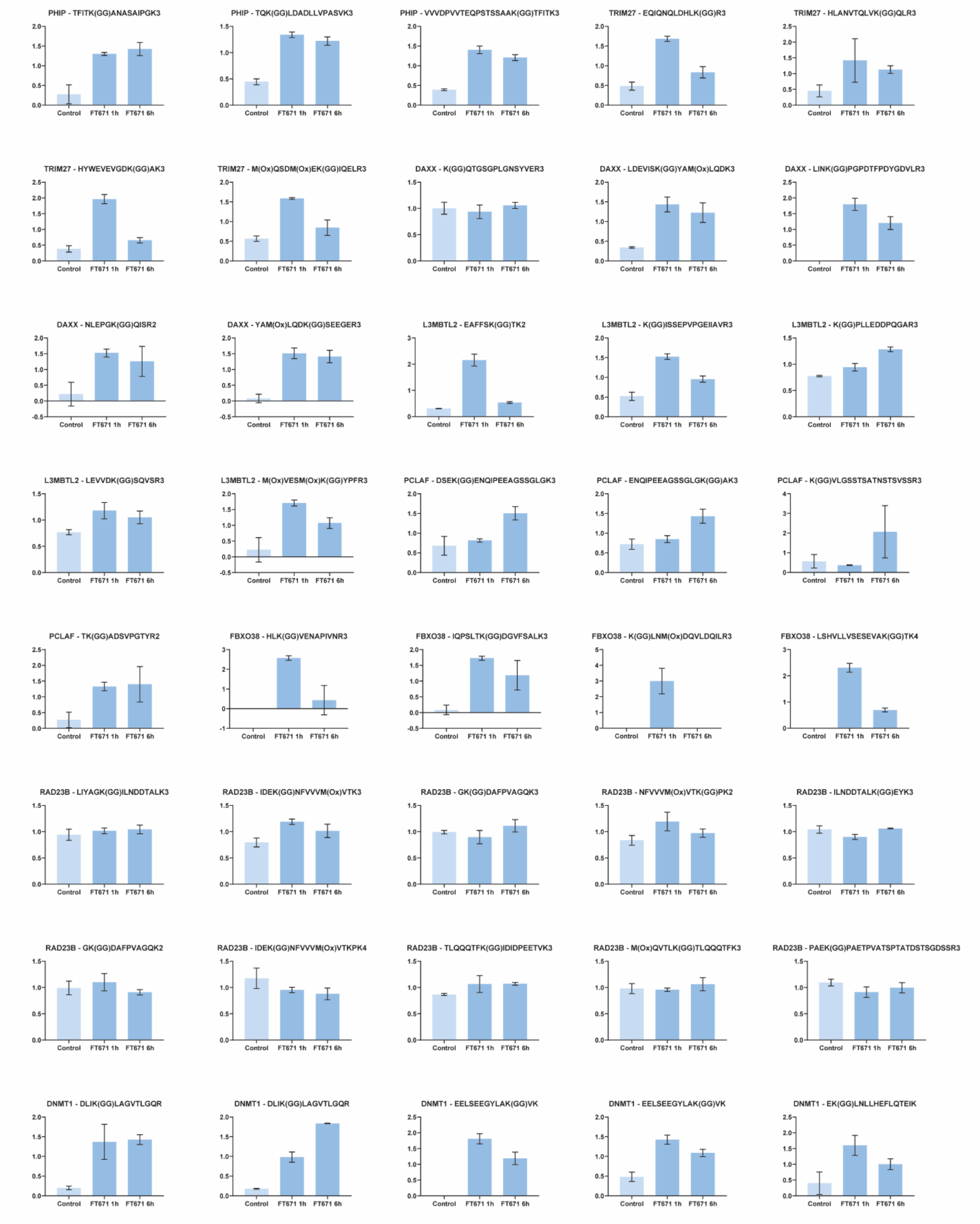
PHIP - TFITK(GG)ANASAIPGK3
PHIP - TQK(GG)LDADLLVPASVK3
PHIP - VVVDPVVTEQPSTSSAAK(GG)TFITK3
TRIM27 - EQIQNQLDHLK(GG)R3
TRIM27 - HLANVTQLVK(GG)QLR3
TRIM27 - HYWEVEVGDK(GG)AK3
TRIM27 - M(Ox)QSDM(Ox)EK(GG)IQELR3
DAXX - K(GG)QTGSGPLGNSYVER3
DAXX - LDEVISK(GG)YAM(Ox)LQDK3
DAXX - LINK(GG)PGPDTFPDYGDVLR3
DAXX - NLEPGK(GG)QISR2
DAXX - YAM(Ox)LQDK(GG)SEEGER3
L3MBTL2 - EAFFSK(GG)TK2
L3MBTL2 - K(GG)ISSEPVPGEIIAVR3
L3MBTL2 - K(GG)PLLEDDPQGAR3
L3MBTL2 - LEVVDK(GG)SQVSR3
L3MBTL2 - M(Ox)VESM(Ox)K(GG)YPFR3
PCLAF - DSEK(GG)ENQIPEEAGSSGLGK3
PCLAF - ENQIPEEAGSSGLGK(GG)AK3
PCLAF - K(GG)VLGSSTSATNSTSVSSR3
PCLAF - TK(GG)ADSVPGTYR2
FBXO38 - HLK(GG)VENAPIVNR3
FBXO38 - IQPSLTK(GG)DGVFSALK3
FBXO38 - K(GG)LNM(Ox)DQVLDQILR3
FBXO38 - LSHVLLVSESEVAK(GG)TK4
RAD23B - LIYAGK(GG)ILNDDTALK3
RAD23B - IDEK(GG)NFVVVM(Ox)VTK3
RAD23B - GK(GG)DAFPVAGQK3
RAD23B - NFVVVM(Ox)VTK(GG)PK2
RAD23B - ILNDDTALK(GG)EYK3
RAD23B - GK(GG)DAFPVAGQK2
RAD23B - IDEK(GG)NFVVVM(Ox)VTKPK4
RAD23B - TLQQQTFK(GG)IDIDPEETVK3
RAD23B - M(Ox)QVTLK(GG)TLQQQTFK3
RAD23B - PAEK(GG)PAETPVATSPTATDSTSGDSSR3
DNMT1 - DLIK(GG)LAGVTLGQR
DNMT1 - DLIK(GG)LAGVTLGQR
DNMT1 - EELSEEGYLAK(GG)VK
DNMT1 - EELSEEGYLAK(GG)VK
DNMT1 - EK(GG)LNLLHEFLQTEIK
Control
FT671 1h
FT671 6h

**Fig. S4.**

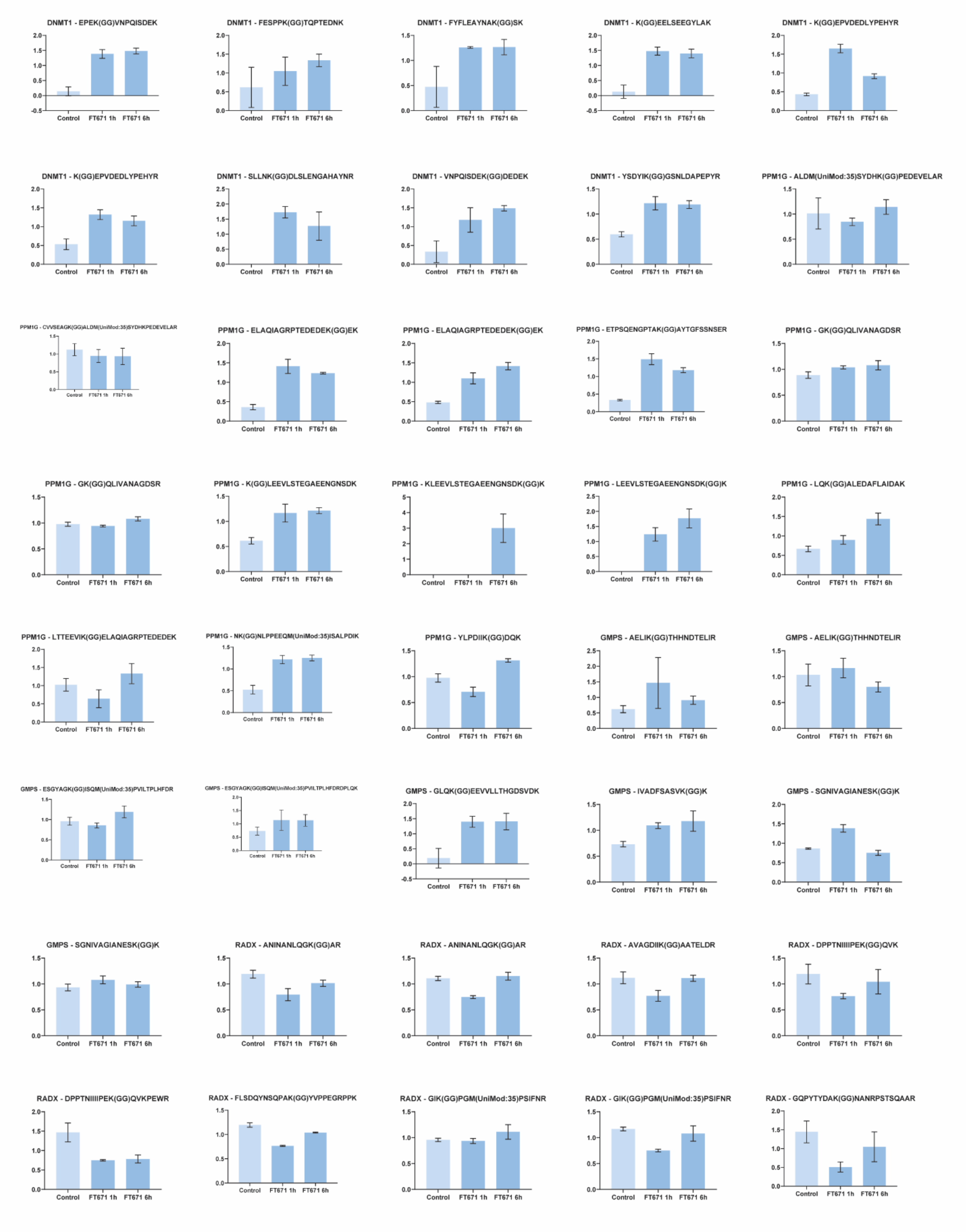
DNMT1 - EPEK(GG)VNPQISDEK
DNMT1 - FESPPK(GG)TQPTEDNK
DNMT1 - FYFLEAYNAK(GG)SK
DNMT1 - K(GG)EELSEEGYLAK
DNMT1 - K(GG)EPVDEDLYPEHYR
DNMT1 - K(GG)EPVDEDLYPEHYR
DNMT1 - SLLNK(GG)DLSLENGAHAYNR
DNMT1 - VNPQISDEK(GG)DEDEK
DNMT1 - YSDYIK(GG)GSNLDAPEPYR
PPM1G - ALDM(UniMod:35)SYDHK(GG)PEDEVELAR
PPM1G - CVVSEAGK(GG)ALDM(UniMod:35)SYDHKPEDEVELAR
PPM1G - ELAQIAGRPTEDEDEK(GG)EK
PPM1G - ELAQIAGRPTEDEDEK(GG)EK
PPM1G - ETPSQENGPTAK(GG)AYTGFSSNSER
PPM1G - GK(GG)QLIVANAGDSR
PPM1G - GK(GG)QLIVANAGDSR
PPM1G - K(GG)LEEVLSTEGAEENGNSDK
PPM1G - KLEEVLSTEGAEENGNSDK(GG)K
PPM1G - LEEVLSTEGAEENGNSDK(GG)K
PPM1G - LQK(GG)ALEDAFLAIDAK
PPM1G - LTTEEVIK(GG)ELAQIAGRPTEDEDEK
PPM1G - NK(GG)NLPPEEQM(UniMod:35)ISALPDIK
PPM1G - YLPDIIK(GG)DQK
GMPS - AELIK(GG)THHNDTELIR
GMPS - AELIK(GG)THHNDTELIR
GMPS - ESGYAGK(GG)ISQM(UniMod:35)PVILTPLHFDR
GMPS - ESGYAGK(GG)ISQM(UniMod:35)PVILTPLHFDRDPLQK
GMPS - GLQK(GG)EEVVLLTHGDSVDK
GMPS - IVADFSASVK(GG)K
GMPS - SGNIVAGIANESK(GG)K
GMPS - SGNIVAGIANESK(GG)K
RADX - ANINANLQGK(GG)AR
RADX - ANINANLQGK(GG)AR
RADX - AVAGDIIK(GG)AATELDR
RADX - DPPTNIIIIPEK(GG)QVK
RADX - DPPTNIIIIPEK(GG)QVKPEWR
RADX - FLSDQYNSQPAK(GG)YVPPEGRPPK
RADX - GIK(GG)PGM(UniMod:35)PSIFNR
RADX - GIK(GG)PGM(UniMod:35)PSIFNR
RADX - GQPYTYDAK(GG)NANRPSTSQAAR

**Fig. S4.**

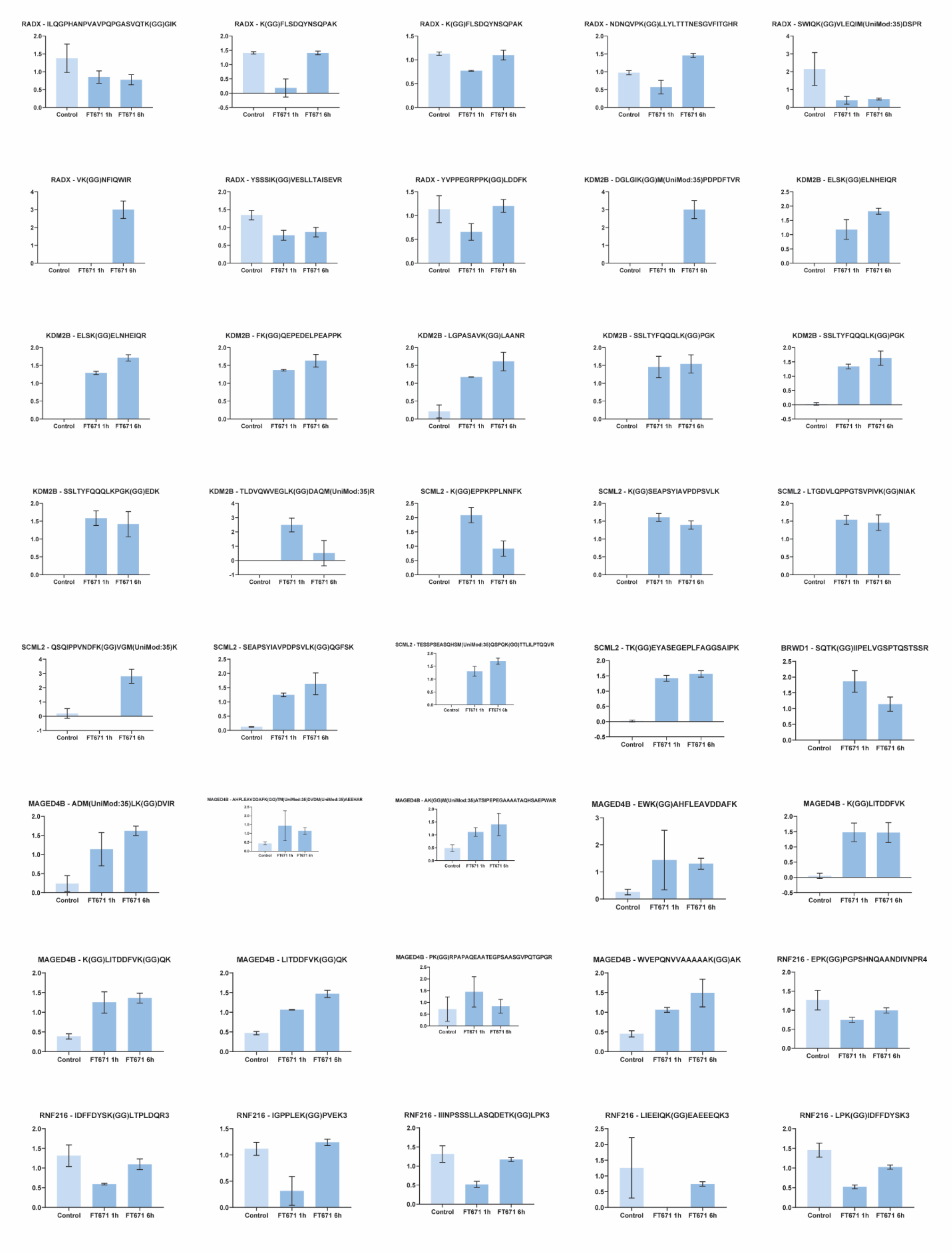
RADX - ILQGPHANPVAVPQPGASVQTK(GG)GIK
RADX - K(GG)FLSDQYNSQPAK
RADX - K(GG)FLSDQYNSQPAK
RADX - NDNQVPK(GG)LLYLTTTNESGVFITGHR
RADX - SWIQK(GG)VLEQIM(UniMod:35)DSPR
RADX - VK(GG)NFIQWIR
RADX - YSSSIK(GG)VESLLTAISEVR
RADX - YVPPEGRPPK(GG)LDDFK
KDM2B - DGLGIK(GG)M(UniMod:35)PDPDFTVR
KDM2B - ELSK(GG)ELNHEIQR
KDM2B - ELSK(GG)ELNHEIQR
KDM2B - FK(GG)QEPEDELPEAPPK
KDM2B - LGPASAVK(GG)LAANR
KDM2B - SSLTYFQQQLK(GG)PGK
KDM2B - SSLTYFQQQLK(GG)PGK
KDM2B - SSLTYFQQQLKPGK(GG)EDK
KDM2B - TLDVQWVEGLK(GG)DAQM(UniMod:35)R
SCML2 - K(GG)EPPKPPLNNFK
SCML2 - K(GG)SEAPSYIAVPDPSVLK
SCML2 - LTGDVLQPPGTSVPIVK(GG)NIAK
SCML2 - QSQIPPVNDFK(GG)VGM(UniMod:35)K
SCML2 - SEAPSYIAVPDPSVLK(GG)QGFSK
SCML2 - TESSPSEASQHSM(UniMod:35)QSPQK(GG)TTLILPTQQVR
SCML2 - TK(GG)EYASEGEPLFAGGSAIPK
BRWD1 - SQTK(GG)IIPELVGSPTQSTSSR
MAGED4B - ADM(UniMod:35)LK(GG)DVIR
MAGED4B - AHFLEAVDDAFK(GG)TM(UniMod:35)DVDM(UniMod:35)AEEHAR
MAGED4B - AK(GG)M(UniMod:35)ATSIPEPEGAAAATAQHSAEPWAR
MAGED4B - EWK(GG)AHFLEAVDDAFK
MAGED4B - K(GG)LITDDFVK
MAGED4B - K(GG)LITDDFVK(GG)QK
MAGED4B - LITDDFVK(GG)QK
MAGED4B - PK(GG)RPAPAQEAATEGPSAASGVPQTGPGR
MAGED4B - WVEPQNVVAAAAAK(GG)AK
RNF216 - EPK(GG)PGPSHNQAANDIVNPR4
RNF216 - IDFFDYSK(GG)LTPLDQR3
RNF216 - IGPPLEK(GG)PVEK3
RNF216 - IIINPSSSLLASQDETK(GG)LPK3
RNF216 - LIEEIQK(GG)EAEEEQK3
RNF216 - LPK(GG)IDFFDYSK3
Control
FT671 1h
FT671 6h

**Fig. S4.**

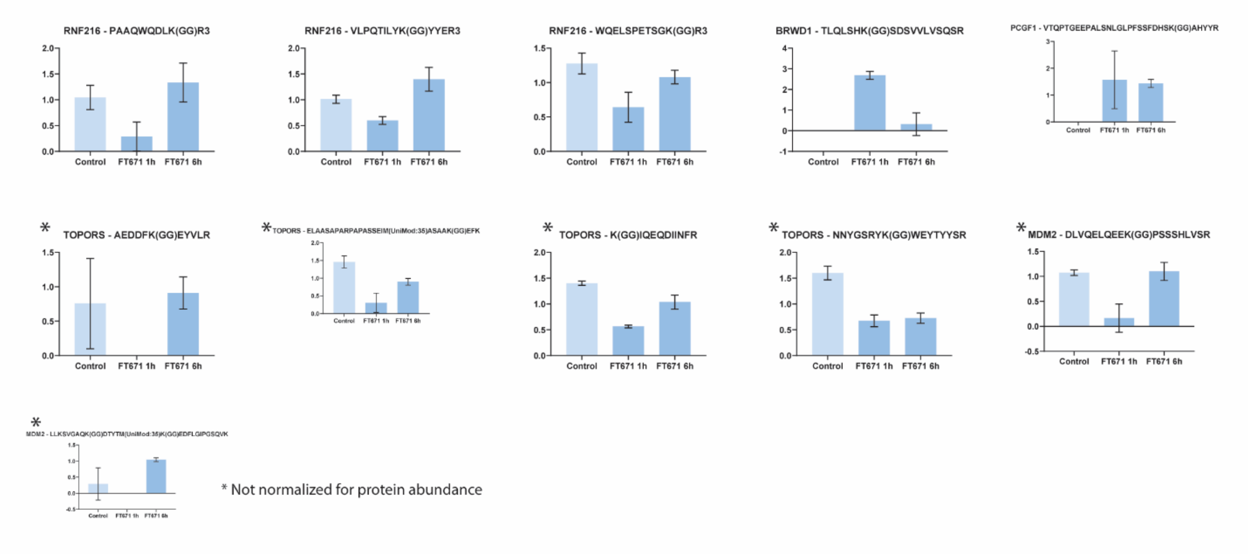


### Fig. S4. Impact of USP7i on the ubiquitylation of high-confidence substrates.

Bar graphs representing scaled relative MS intensities representing K-GG peptide abundances that are normalized for overall protein abundance (except those indicated with a "*"), in HEK293T cells cultured in the presence of 7 mM DMSO (6 h), or 10 µM FT671 for 1 or 6 h. See also Supplemental Table S7 and 9.
